# Identification of key genes and enriched biological processes in diabetes-associated traumatic brain injury through weighted gene correlation network and bioinformatics analysis

**DOI:** 10.1101/2024.10.12.615673

**Authors:** Umar Faruk Saidu, Ibrahim Bulama

**Affiliations:** Department of Biochemistry and Molecular Biology, Faculty of Life Sciences, Usmanu Danfodiyo University, Sokoto, Nigeria; Department of Veterinary Physiology and Biochemistry, Faculty of Veterinary Medicine, University of Maiduguri, Maiduguri, Nigeria

**Keywords:** Traumatic brain injury, Type 1 diabetes, Weighted gene co-expression network analysis, Cmklr1, Mgst1, Plin2, RNA-binding proteins, Neutrophil degranulation, Lipid droplet dysregulation

## Abstract

Traumatic brain injury (TBI) is a significant cause of morbidity and mortality worldwide, with long-term neurological and psychological impacts. Recent studies have indicated that diabetes (T1DM) exacerbates the outcomes of TBI, leading to more severe cognitive deficits and increased risk of complications. This study investigated the underlying molecular mechanisms and potential therapeutic targets for T1DM-associated TBI. Four mRNA datasets (GSE4745, GSE125451, GSE173975, and GSE80174) downloaded from GEO repository were used in this study. Using limma, a total of 284 differentially expressed genes (DEGs) were identified in T1DM, of which 11 were upregulated and 9 were downregulated. GSEA showed that these DEGs were significantly enriched in cell communication, lipid metabolic process, and PPAR signaling. A total of 584 DEGs were identified in TBI, of which 186 were upregulated and 9 were downregulated. GSEA showed that these DEGs were mainly enriched in immune response-regulating signaling pathway. WGCNA identified 122 significant genes in TIDM-related modules and 368 significant genes in TBI-related module. GO and KEGG enrichment analysis showed that T1DM module genes were significantly correlated with lipid metabolic process and ribosome biogenesis, while TBI module genes were significantly correlated with inflammation and immune response, including leukocyte mediated immunity, lymphocyte mediated immunity, and cytokine mediated receptor activity. PPI network analysis of T1DM module genes identified 20 hub genes, including 14 ribosomal genes: Rpl23, Rps3a, Rps6, Rpl5, Rpl17, Rps24, Rpl23a, Rps4x, Rpl9, Rps15a14, Rpl30, Rpl31, Rps25, and Rps27a-2. The hub genes were primarily related to ribosome biogenesis and RNA post-transcriptional regulation. PPI network analysis of TBI module genes identified 20 hub genes: Ptprc, Tp53, Stat1, Stat3, Tyrobp, Itgad, Csf1r, Itgb2, Rac2, Icam1, Myd88, Cd44, Vav1, Aif1, C1qa, Laptm5, B2m, Fcer1g, and Lyn. The hub genes were primarily related to inflammatory mediators and immune response. Based on the overlap of T1DM module genes and TBI module genes, Cmklr1, Mgst1, and Plin2, were identified as key genes of T1DM-associated TBI. Functional enrichment analysis showed that they were primarily enriched in the cellular response to hydroperoxide, cytokine-mediated receptor signaling activity, regulation of sequestering of triglyceride, negative regulation of IL-12 production, and positive regulation of macrophage chemotaxis. Using Reactome, Cmklr1, Mgst1, and Plin2 were related to cytokine signaling activity, neutrophil degranulation, and lipid storage, respectively. We concluded that Lipid droplet dysregulation and Neuroinflammation are the potential molecular mechanisms of T1DM-associated TBI. Cmklr1, Mgst1, and Plin2 are positively correlated with T1DM-associated TBI and may be important biomarkers and potential treatment targets for diabetic TBI.

## INTRODUCTION

Traumatic brain injury (TBI) is a significant cause of morbidity and mortality worldwide, with long-term neurological and psychological impacts. An estimated 1.7 million Americans get TBI, costing the nation’s healthcare systems around $76.5 billion a year [1]. The presence of comorbid conditions such as diabetes mellitus exacerbates the outcomes of TBI, leading to more severe cognitive deficits and increased risk of complications [2]. Diabetes, characterized by chronic hyperglycemia, affects various cellular and molecular pathways, which can influence the response to TBI [2].

The interplay between type 1 diabetes (T1DM) and TBI is complex, involving alterations in metabolic pathways, inflammatory responses, and neurovascular integrity [2]. Recent studies have indicated that persistent hyperglycemia can exacerbate the damaging effects of TBI by promoting stress-induced hyperglycemia, oxidative stress, inflammation, pituitary and hypothalamic dysfunction [2, 3, 4]. However, the precise molecular mechanisms underlying these interactions are yet to be explored. By using high-throughput gene expression profiling and bioinformatics analysis to detect changes in gene expression of T1DM and TBI, it is possible to gain insight into the molecular alterations brought about by the combined effects when they coexist.

Weighted gene co-expression analysis (WGCNA) is a systems biology technique one can use to create gene correlation networks and pinpoint significant gene modules linked to particular diseases or biological functions. It calculates gene correlation using gene expression data and groups highly correlated genes into modules to identify gene regulatory networks and important regulatory genes [5]. WGCNA identifies gene modules using unsupervised clustering, i.e., without the use of a priori-defined gene sets [5]. Previous studies have used WGCNA to explore gene expression data, identify gene modules linked to disease pathology, and perform functional enrichment analysis to understand the molecular mechanisms of diseases [6].

This work aimed to identify key genes and enriched biological processes of T1DM-associated TBI through the identification of the shared molecular mechanisms of T1DM and TBI. We downloaded four Rattus norvegicus microarray data of T1DM and TBI from the GEO database. We used WGCNA and bioinformatics analysis to explore the differentially coexpressed genes and signaling pathways in T1DM-associated TBI. GSEA, Metascape, GO, and KEGG functional enrichment analysis were used to understand the biological functions of identified key genes. Protein-protein interaction (PPI) network analysis was used to identify hub genes. We identified the shared genes that might be correlated with the pathophysiology of T1DM-associated TBI based on the overlap of TIDM-related module genes and TBI-related module genes. Finally, Reactome was used to further understand the biological reactions of the shared genes through interactive pathways diagram. The shared genes could serve as novel biomarkers and promising therapeutic targets for T1DM-associated TBI.

## METHODS

### Data Acquisition

From the Gene Expression Omnibus (GEO) database, we downloaded four Rattus norvegicus mRNA data related to T1DM and TBI. There were two datasets related to T1DM (GSE4745 and GSE125451) and two datasets related to TBI (GSE173975 and GSE80174). The following criteria were used for inclusion: (1) The datasets must contain raw mRNA data from Rattus norvegicus; (2) The datasets must contain disease and healthy control samples; (3) The datasets preferably should come from the same tissue: T1DM datasets were from the heart and TBI datasets were from the hippocampus; (4) The disease conditions ideally should be induced using the same model: T1DM was induced using Streptozotocin model and TBI was induced using lateral fluid percussion model; (5) The combined dataset for each of T1DM and TBI were greater than 20 samples; this increases the robustness of the WGCNA analysis.

### Data Assembly and Preprocessing

The raw counts were normalized and converted to cpm values and log transformed using the TMM-normalization method in the “edgeR” package. The two normalized diabetes datasets were then combined together to form a single expression data. The two normalized TBI datasets were also combined together to form a single expression data. Then, the combined expression data were batch corrected using ComBat method in the “sva” package. Finally, genes with low expression values (sum across all samples) below 15 were filtered out from the expression data.

### Differential Gene Expression Analysis

We used the “limma” package to screen for differentially expressed genes (DEGs) in T1DM and TBI expression data by comparing the disease and control groups. The threshold criteria set was a P-value less than 0.05, and absolute log fold change greater than 0.25. The R packages “ComplexHeatmap” and “EnhancedVolcano” were used for plotting the heatmap and volcano plot of the DEGs.

### Gene Set Enrichment Analysis

One popular technique for genomic analysis is “gene set enrichment analysis” (GSEA), which involves querying an input gene list against libraries of annotated gene sets to find gene sets that significantly overlap the input genes [7]. Metascape is a popular gene annotation search engine with a sizable library of gene sets for conducting GSEA analyses [8]. GSEA, and Metascape available at https://metascape.org/gp/index.html#/main/step1 were used to identify significantly enriched biological processes and pathways between the disease and healthy controls. The DEGs with p-value < 0.05 were preranked based on logFC values and used for GSEA.

### Weighted Gene Co-expression Network Analysis (WGCNA)

Applications of correlation networks in bioinformatics are growing in popularity. WGCNA is an algorithm for finding co-expressed gene modules, calculating module membership degrees, correlating modules with clinical traits, and summarizing such clusters using an intramodular hub gene or the module eigengene [9]. The gene hierarchical clustering dendrogram was constructed in order to detect co-expression modules. Spearman correlation analysis and hierarchical clustering of genes were used to determine the module eigengenes (MEs) and the association between traits and MEs in order to find clinically correlated modules.

### GO and KEGG Enrichment Analysis of Genes in Disease-related Key Modules

The R package “clusterProfiler” tool was used to conduct GO and KEGG analysis for genes in significant modules related to T1DM and TBI. The three separate ontologies that were constructed are molecular function (MF), cellular component (CC), and biological process (BP). Kyoto Encyclopedia of Genes and Genomes (KEGG) enrichment analysis was used for pathway analysis.

### Protein Network Analysis and Identification of Hub Genes of T1DM and TBI

The protein–protein interaction (PPI) networks were constructed using the STRING website (https://string-db.org/) (version 11.5) [10]. We used STRING to create PPI networks of WGCNA significant module genes with an interaction score above 0.4. We entered the gene symbols into the website and then picked multiple proteins, using the gene symbols as a list of names and Rattus norvegicus as the organism. The networks were exported to Cytoscape software and CytoHubba (version 3.9.1) plug-in was used to identify the hub genes of T1DM and TBI.

### Identification of Shared Genes of T1DM-associated TBI

After performing WGCNA on expression data to identify disease-related modules, the shared genes were identified by intersecting T1DM-related significant modules and TBI-related significant modules to find genes of T1DM-associated TBI.

### Functional Enrichment Analysis of Shared Genes of T1DM-associated TBI

We used “clusterProfiler” to identify highly enriched terms (P value ≤ 0.01) for the GO keywords— biological process (BP), cellular component (CC), and molecular function (MF) of the shared genes. Furthermore, using the Reactome database (https://reactome.org), we identified and visualized the interactive biological pathways of the shared genes.

## RESULTS

### Identification of DEGs in T1DM and TBI

The four GEO datasets that were used in this study are summarized in **Table S1**, and the flowchart of the bioinformatics work flow is shown in **Fig. 1**. Each sample from T1DM and TBI datasets were normalized using TMM-normalization method (cpm and log transformed) in “edgeR” package. The batch effects were corrected using the combat function in “sva” package (**Fig. 2A and B**). We used “limma” R package to find genes that were differentially expressed in T1DM and TBI between the disease and healthy controls. A total of 286 DEGs were identified in T1DM, and 11 genes were upregulated and 9 genes were downregulated. The DEGs were sorted based on p-values and the top 15 T1DM DEGs were visualized using a volcano plot. We then sorted the DEGs based on logFC values, and a clustered heatmap was used to visualize the top 50 DEGs, which included 25 upregulated DEGs and 25 downregulated DEGs (**Fig. 3A and B**). A total of 584 DEGs were identified in TBI, and 186 genes were upregulated and 9 genes were downregulated. Similarly, a volcano plot was used to visualize the top 15 TBI DEGs sorted based on p-values, and a clustered heatmap was used to visualize the top 50 DEGs sorted based on logFC values, which included 25 upregulated and 25 downregulated DEGs (**Fig. 3C and D**).

**Fig. 1.**
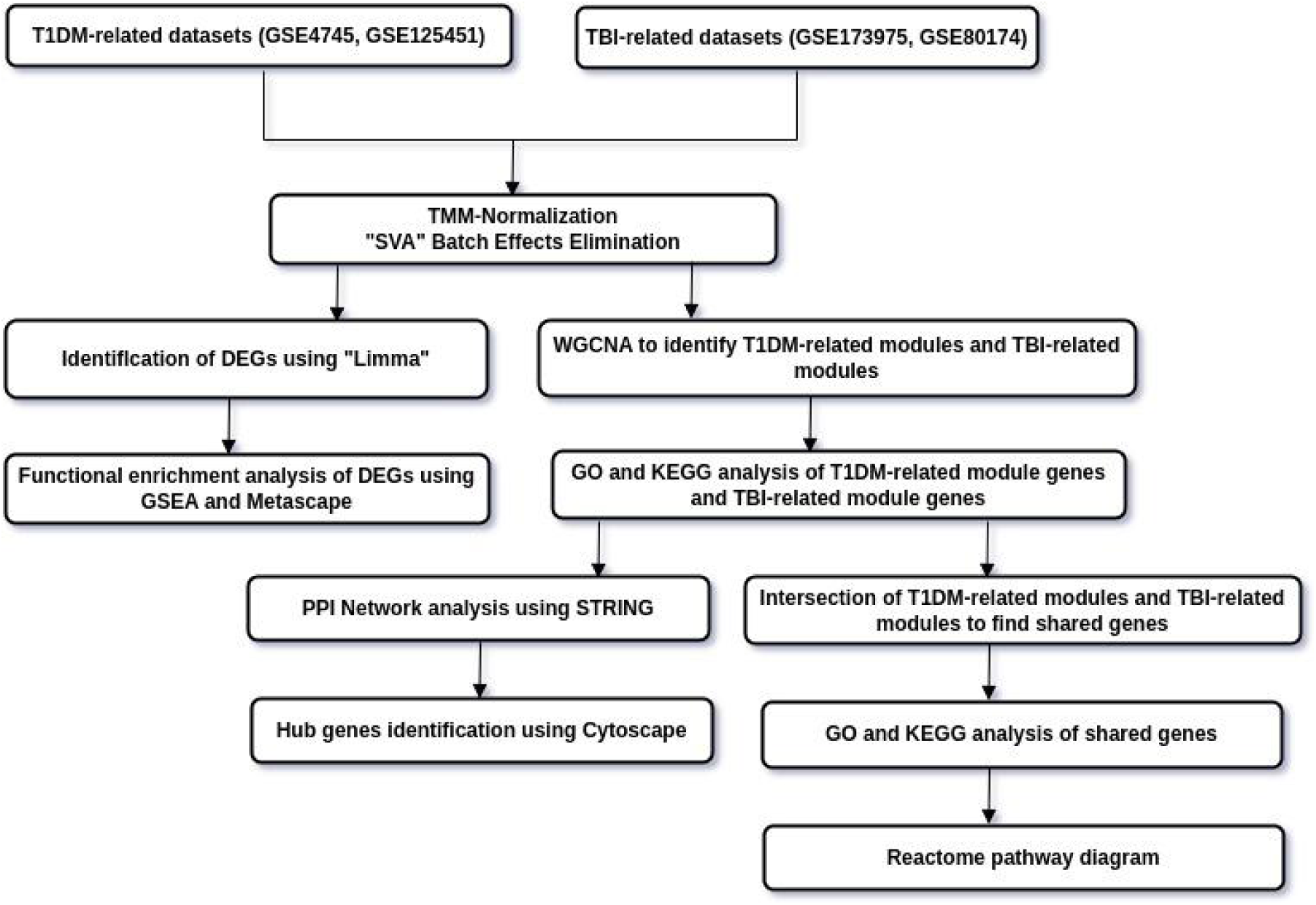
Flowchart of the Bioinformatics work flow. SVA: Surrogate variable analysis; Limma: Linear Models for Microarray and Omics Data; WGCNA: Weighted gene co-expression network analysis; GSEA: Gene set enrichment analysis; GO: Gene Ontology; KEGG: Kyoto Encyclopaedia of Genes and Genomes; PPI: protein–protein interaction; STRING: Search Tool for the Retrieval of Interacting Genes/Proteins.

**Fig. 2.**
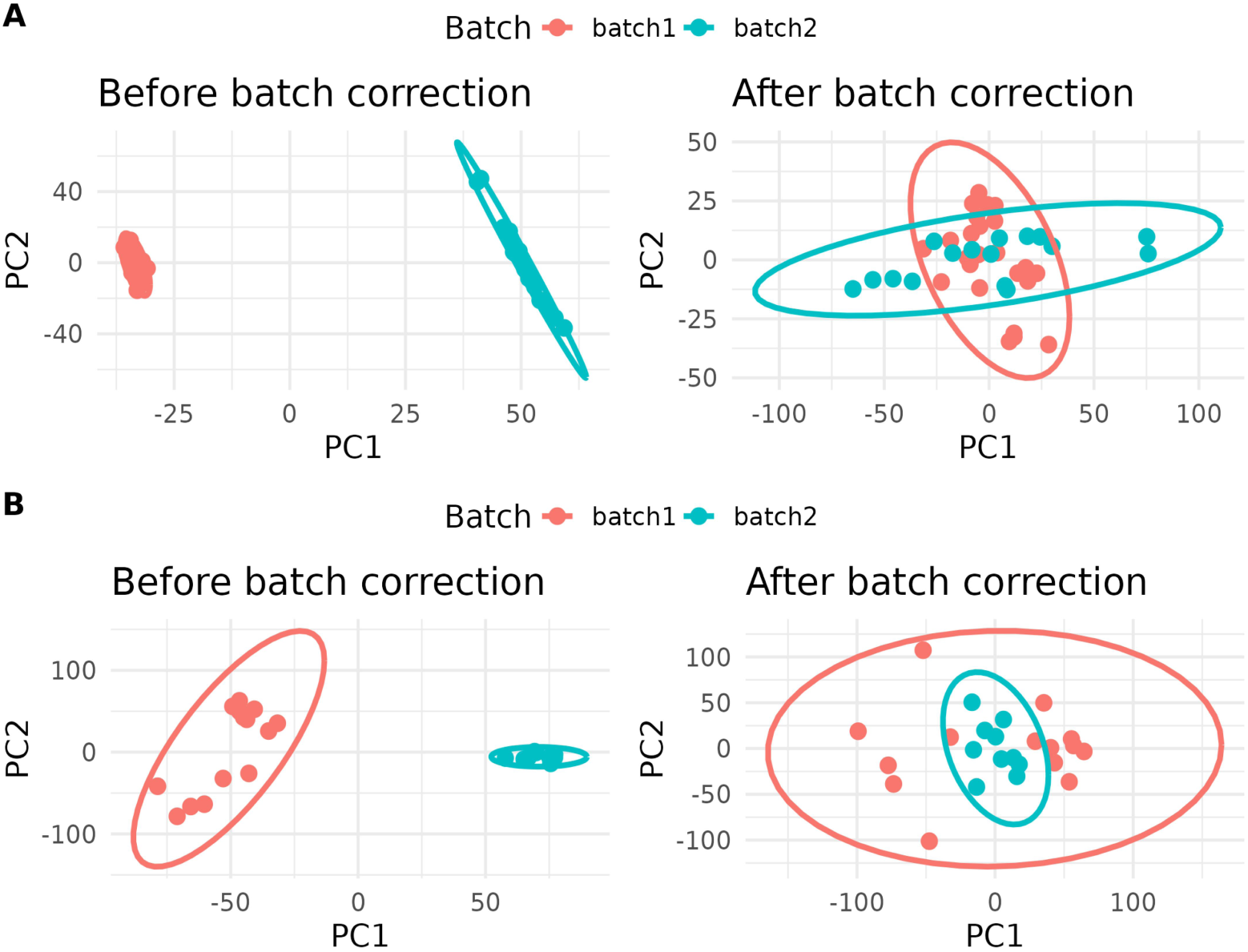
Batch effects elimination of mRNA data using ComBat. (A) Batch correction of T1DM dataset. First frame is before batch correction and second frame is after batch correction. (B) Batch correction of TBI dataset. First frame is before batch correction and second frame is after batch correction.

**Fig. 3.**
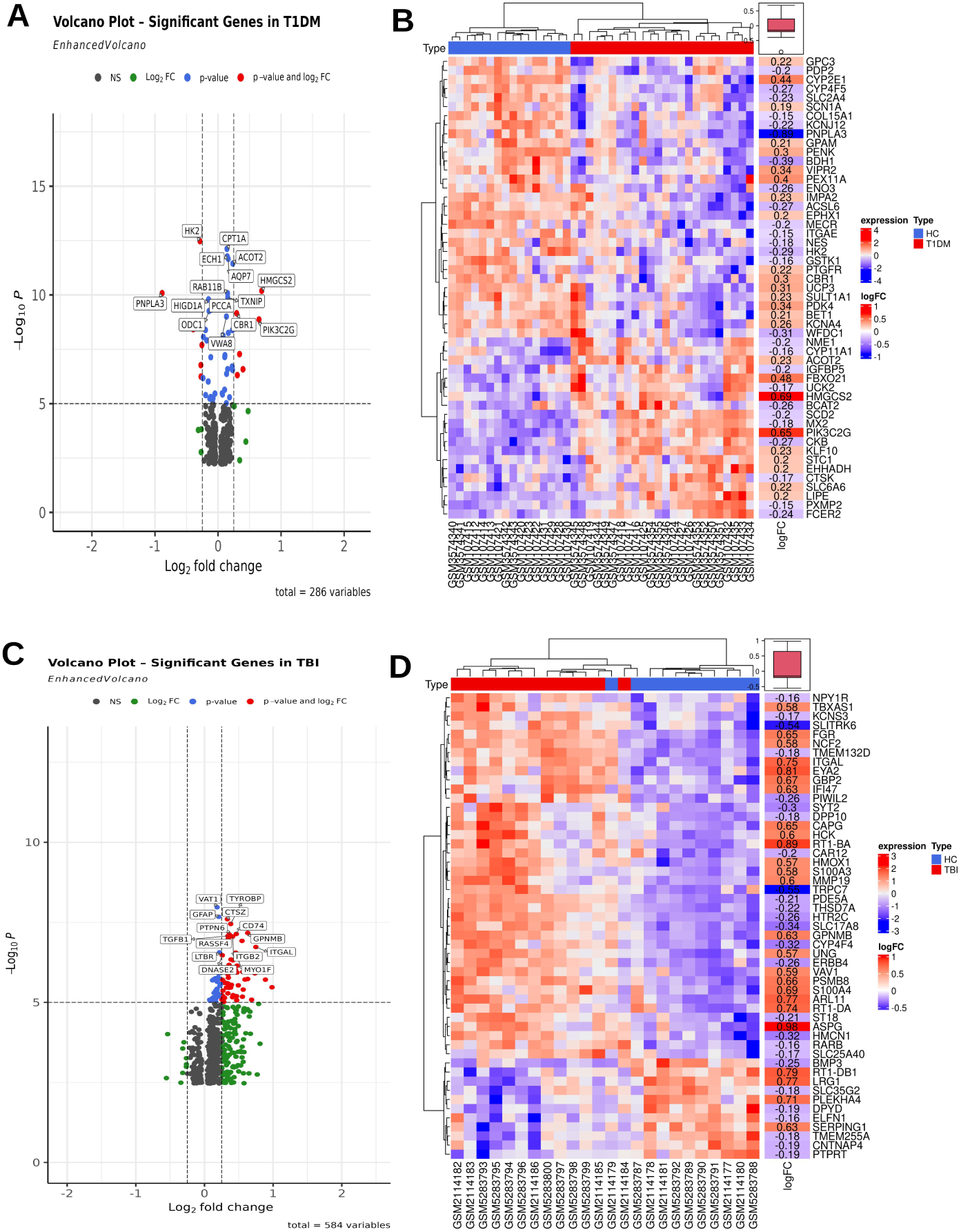
Identification of T1DM and TBI DEGs using Limma. (A) A volcano plot of the top 15 T1DM DEGs. The DEGs were sorted in descending order based on p-values. (B) A clustered heatmap of the top 50 T1DM DEGs. The DEGs were sorted in descending order based on logFC values, including 25 upregulated DEGs and 25 downregulated DEGs. (C) A volcano plot of the top 15 TBI DEGs. The DEGs were sorted in descending order based on p-values. (D) A clustered heatmap of the top 50 TBI DEGs. The DEGs were sorted in descending order based on logFC values, including 25 upregulated DEGs and 25 downregulated DEGs. T1DM – Type 1 diabetes mellitus. TBI – Traumatic brain injury. DEGs – Differentially expressed genes.

### Gene Set Enrichment Analysis (GSEA) of T1DM and TBI

GSEA was used to identify probable molecular processes underlying T1DM and TBI. The GSEA results identified cell communication, response to external stimulus and response to lipid metabolic process as the most enriched biological processes in T1DM. The dot plot showed that these processes were activated in T1DM compared to healthy controls. We have similar enrichment distribution pattern using the ridge plot (**Fig. 4A, B, and C**). Using Metascape to further annotate the DEGs, the most enriched pathways in T1DM were related to metabolism of monocarboxylic acids, the response to xenobiotic stimulus, PPAR signaling pathway, metabolism of lipid (fatty acid metabolism, fatty acid beta-oxidation, and COA carboxylase activity), metabolism of glutathione, and IL-2/Stat5 signaling (**Fig. 4D**). The GSEA results identified immune response and immune system process as the most enriched biological processes in TBI. The dot plot showed that these processes were activated in TBI compared to healthy controls. We used a ridge plot to show the enrichment distribution pattern of these biological processes in TBI (**Fig. 5A, B, and C**). The metascape enrichment analysis showed that the DEGs were primarily associated with immune response-regulating signaling pathways, including innate immune response, response to stress, cell activation, neutrophil activation, and degranulation, positive regulation of tumor necrosis factor production, positive regulation of reactive oxygen species, Th1 and Th2 cell differentiation, Toll-like-receptor signaling pathway, and positive regulation of IL-6 production (**Fig. 5D)**. According to the GSEA results, the commonly enriched biological processes in T1DM and TBI were immune system response, response to external stimulus, and cell activation.

**Fig. 4.**
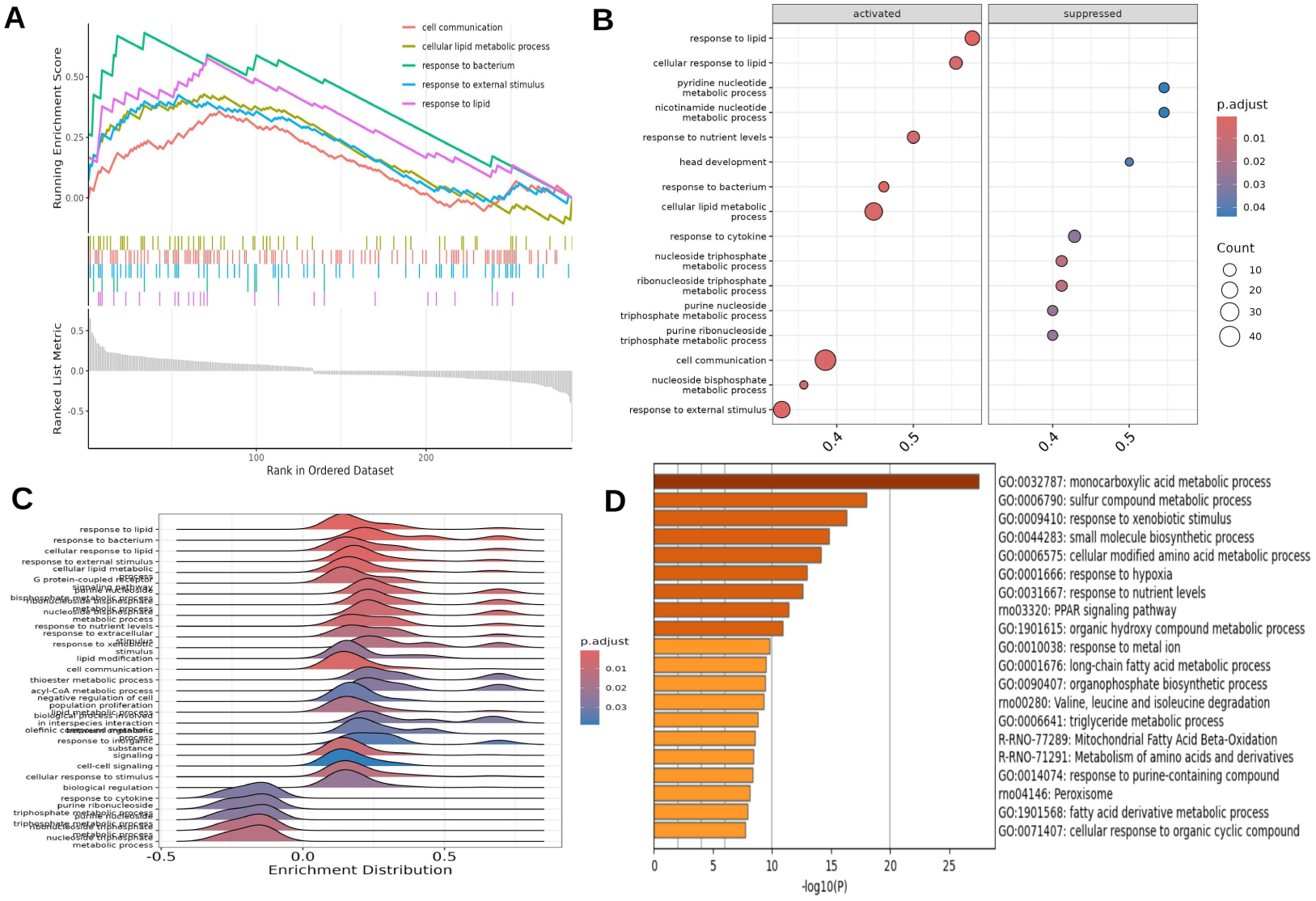
Functional enrichment of T1DM DEGs using GSEA and Metascape. (A) The GSEA of the top 5 enriched biological processes in T1DM gene set. The DEGs were preranked in descending order based on logFC values for GSEA analysis. (B) Dot plot of the top activated and suppressed biological processes in T1DM compared to healthy controls. (C) Ridge plot of the enrichment distribution pattern of the enriched biological processes. (D) The enriched GO terms of the T1DM gene set using Metascape. T1DM – Type 1 diabetes mellitus. DEGs – Differentially expressed genes. GO – Gene ontology.

**Fig. 5.**
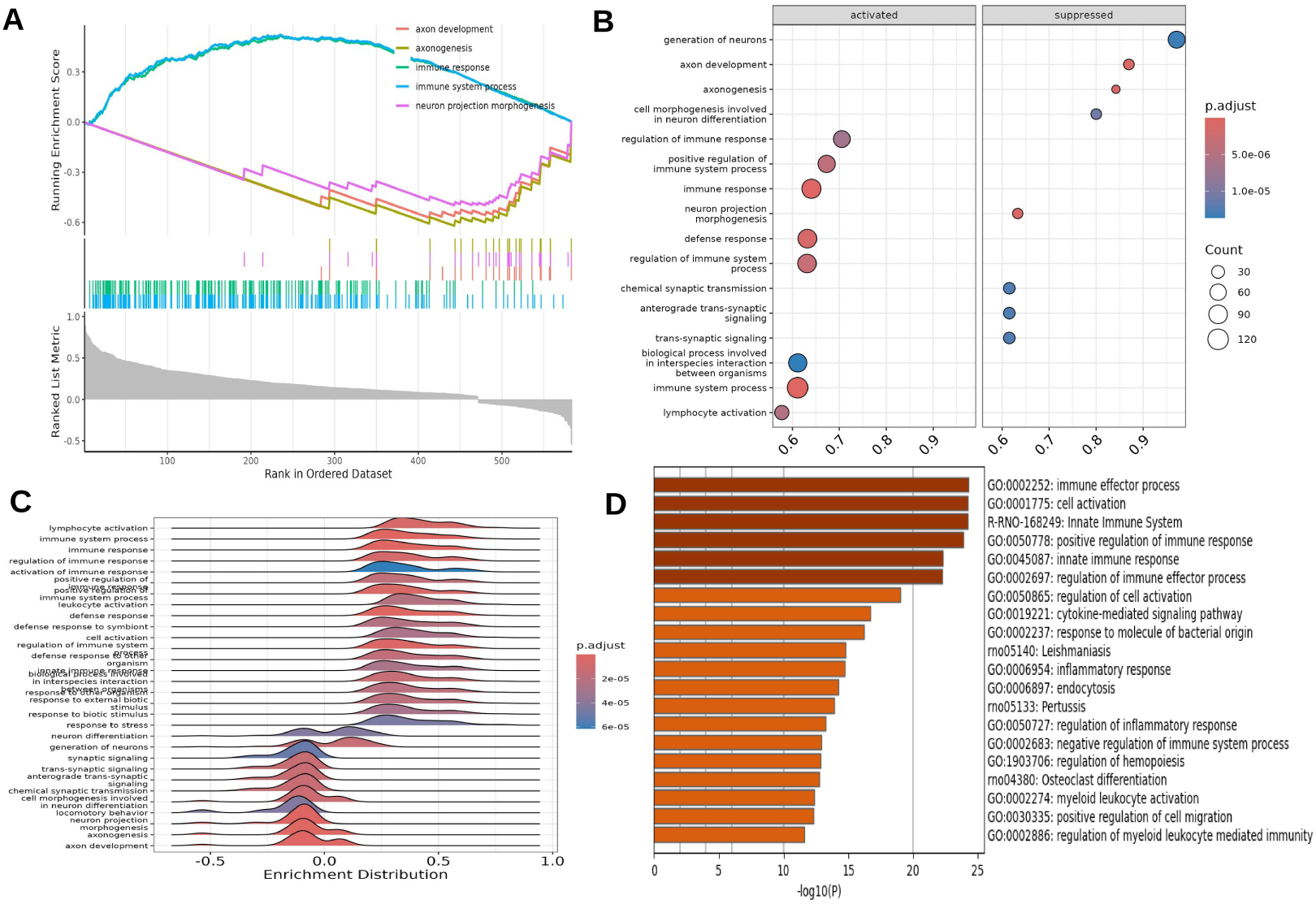
Functional enrichment of TBI DEGs using GSEA and Metascape. (A) The GSEA of the top 5 enriched biological processes in TBI gene set. The DEGs were preranked in descending order based on logFC values for GSEA analysis. (B) Dot plot of the top activated and suppressed biological processes in TBI compared to healthy controls. (C) Ridge plot showing the enrichment distribution pattern of enriched biological processes. (D) The enriched GO terms of the gene set using Metascape. TBI – Traumatic brain injury. DEGs – Differentially expressed genes. GO – Gene ontology.

### Network Construction and Identification of T1DM-related Key Modules

We used WGCNA with TOMType “Signed” network to identify disease-related modules in the T1DM dataset. The network topology analysis was visualized for different soft threshold powers and picking a power near the curve of the plot (scale-free topological index 0.8), a power of 10 was selected for network construction and module identification (**Fig. 6A**). We calculated the correlations between T1DM and healthy controls using hierarchical clustering of genes into distinct modules and spearman correlation analysis (**Fig. 6B**). We effectively identified 10 different modules and the red module (r = 0.80, p = 2.22e-308) and black module (r = 0.46, p = 2.78e-75) were chosen as the significant modules because of their highest correlation with T1DM (**Fig. 6C**), and a total of 122 genes from these modules were selected (GS > 0.25 and MM > 0.8) (**Fig. 6D**). To identify the molecular mechanism underlying T1DM, we performed GO and KEGG functional enrichment analysis of the 122 genes (**Fig. 7A, B, C, and D**). The enriched GO biological processes were the fatty acid metabolic process, response to hypoxia, cellular lipid catabolic process, and sulfur compound metabolic process (Fig. 7A). These genes were mainly enriched in the cytosolic ribosome, ribosomal subunit, peroxisome, and mitochondrial matrix cellular components (**Fig. 7B**). There molecular functions were predominantly in structural constituent of ribosome, rRNA binding, and fatty-acyl-CoA binding activity (**Fig. 7C**). The KEGG pathway analysis identified ribosome biogenesis, peroxisomal pathway, insulin signaling pathway, PPAR signaling pathway, and AMPK signaling pathway to be significantly enriched (**Fig. 7D**).

**Fig. 6.**
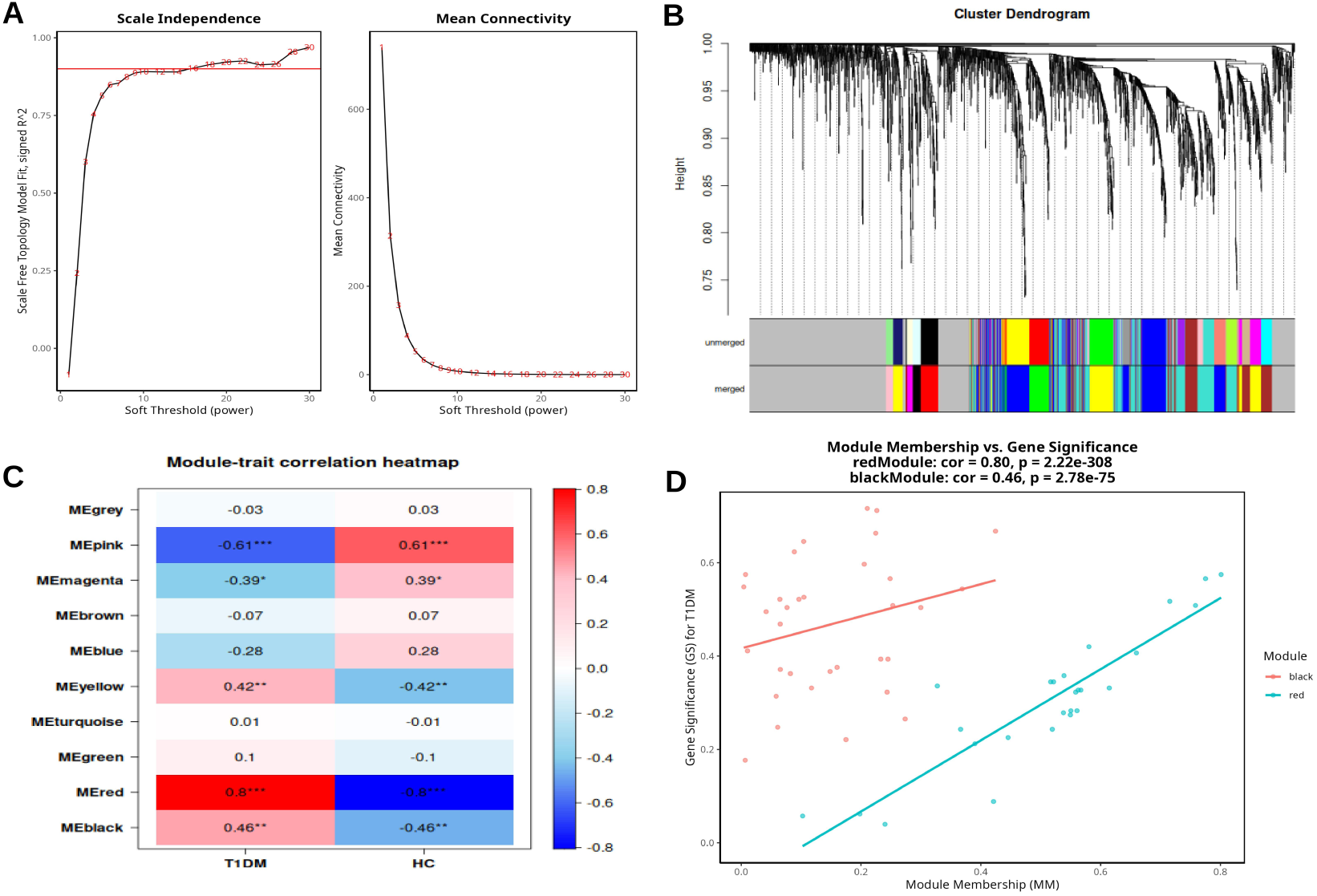
Network construction and identification of T1DM-related key modules. (A) The Scale independence and Mean connectivity for different powers. A power of 10 having highest scale free topology and lowest mean connectivity was chosen for network construction based on the plot. (B) The hierarchical clustering of genes into distinct modules. (C) Module-trait correlation heatmap. Modules were correlated with traits using Spearman correlation. (D) Module Membership vs Gene Significance of the selected significant modules.

**Fig. 7.**
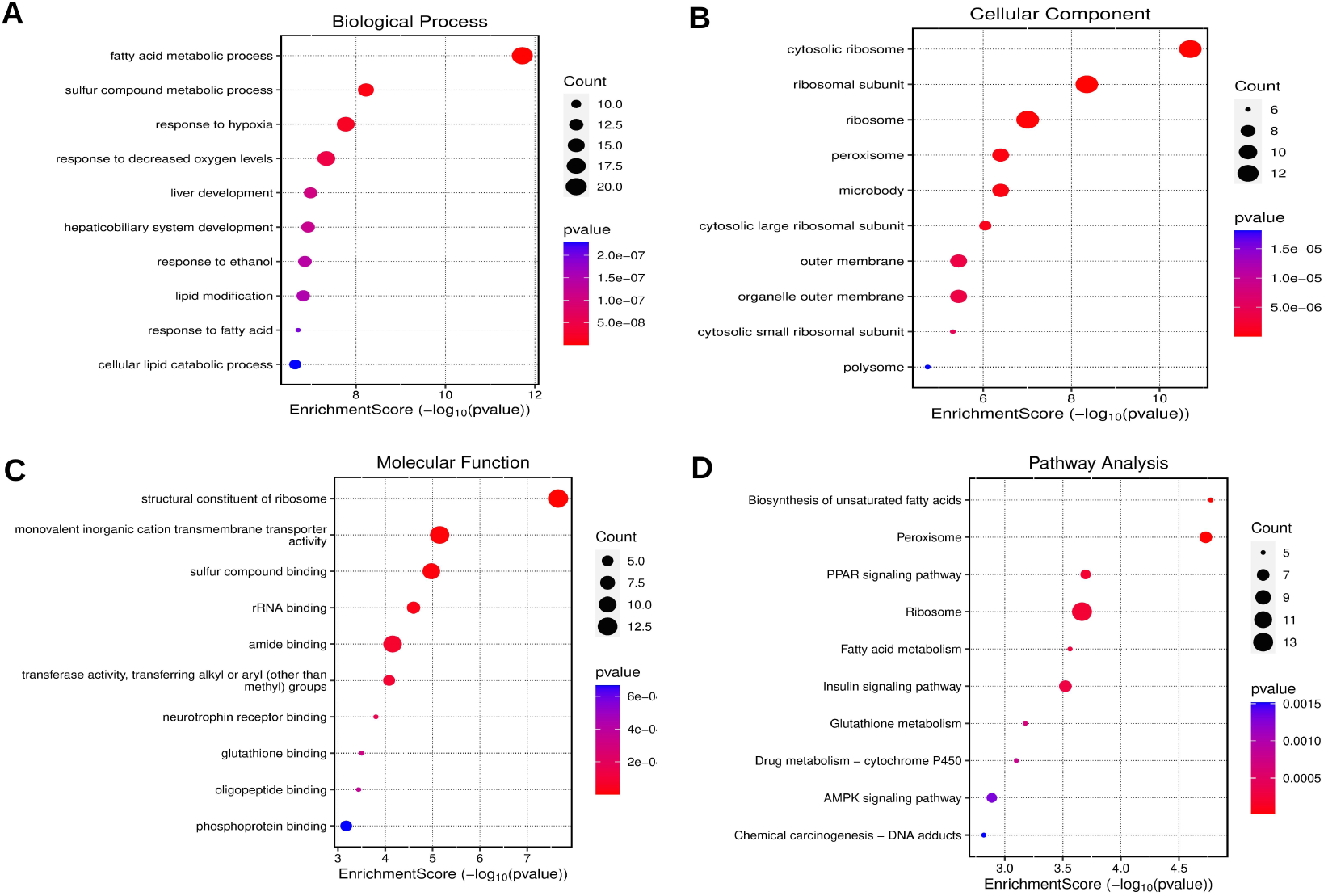
GO and KEGG enrichment analysis of T1DM-related module genes using clusterProfiler. (A) The top enriched Biological process (B) The top enriched Cellular component (C) The top enriched Molecular function (D) The top enriched KEGG pathways.

### Network Construction and Identification of TBI-related Key Modules

We used WGCNA with TOMType “Signed” network to identify disease-related modules in the TBI dataset. The network topology analysis was visualized for different soft threshold powers and picking a power near the curve of the plot (scale-free topological index 0.8), a power of 26 was selected for network construction and module identification (**Fig. 8A**). We calculated the correlations between TBI and healthy controls using hierarchical clustering of genes into distinct modules and spearman correlation analysis (**Fig. 8B and C**). We effectively identified 9 different modules and the yellow module (r = 0.83, p = 2.22e-308) was chosen as the significant module because of its highest correlation with TBI (**Fig. 8C**), and 368 yellow module genes were selected (GS > 0.25 and MM > 0.8) (**Fig. 8D**). The 368 genes were functionally annotated using GO and KEGG analysis (**Fig. 9A, B, C, and D**). The enriched GO biological process was mainly associated with immune response-regulating signaling pathways; including leukocyte-mediated immunity, adaptive immune response, myeloid leukocyte activation, proliferation, and migration, lymphocyte-mediated immunity, and the regulation of cell-cell adhesion (**Fig. 9A**). Regarding cell components, genes were predominantly enriched in lytic vacuole, lysosome, cell leading edge, actin filament, phagocytic vesicle, integrin complex, and protein complex involved in cell adhesion (**Fig. 9B**). The genes were enriched in molecular functions such as SH3 domain binding, cell adhesion molecule binding, integrin binding, actin binding, and immune receptor activity (**Fig. 9C**). The KEGG analysis showed a strong correlation between TBI and the enrichment of chemokine signaling pathway, the NOD-like signaling pathway, Natural killer cell mediated cytotoxicity, Fc gamma R-mediated phagocytosis, lysosomal pathway and the complement cascade/coagulation signaling pathway (**Fig. 9D**).

**Fig. 8.**
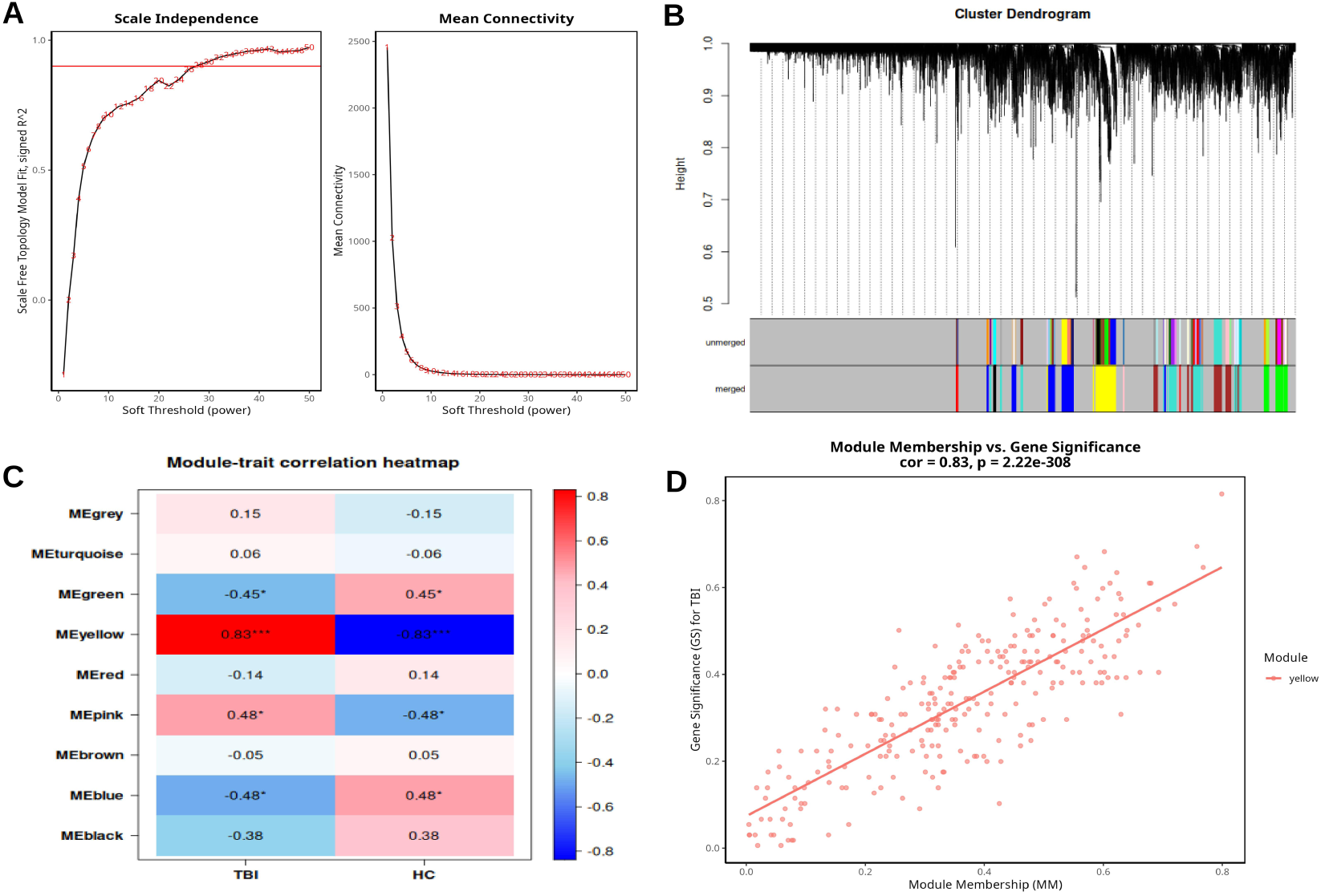
Network construction and identification of TBI-related key modules. (A) The Scale independence and Mean connectivity for different powers. A power of 26 having highest scale free topology and lowest mean connectivity was chosen for network construction based on the plot. (B) The hierarchical clustering of genes into distinct modules. (C) Module-trait correlation heatmap. Modules were correlated with traits using Spearman correlation. (D) Module Membership vs Gene Significance of the selected significant module.

**Fig. 9.**
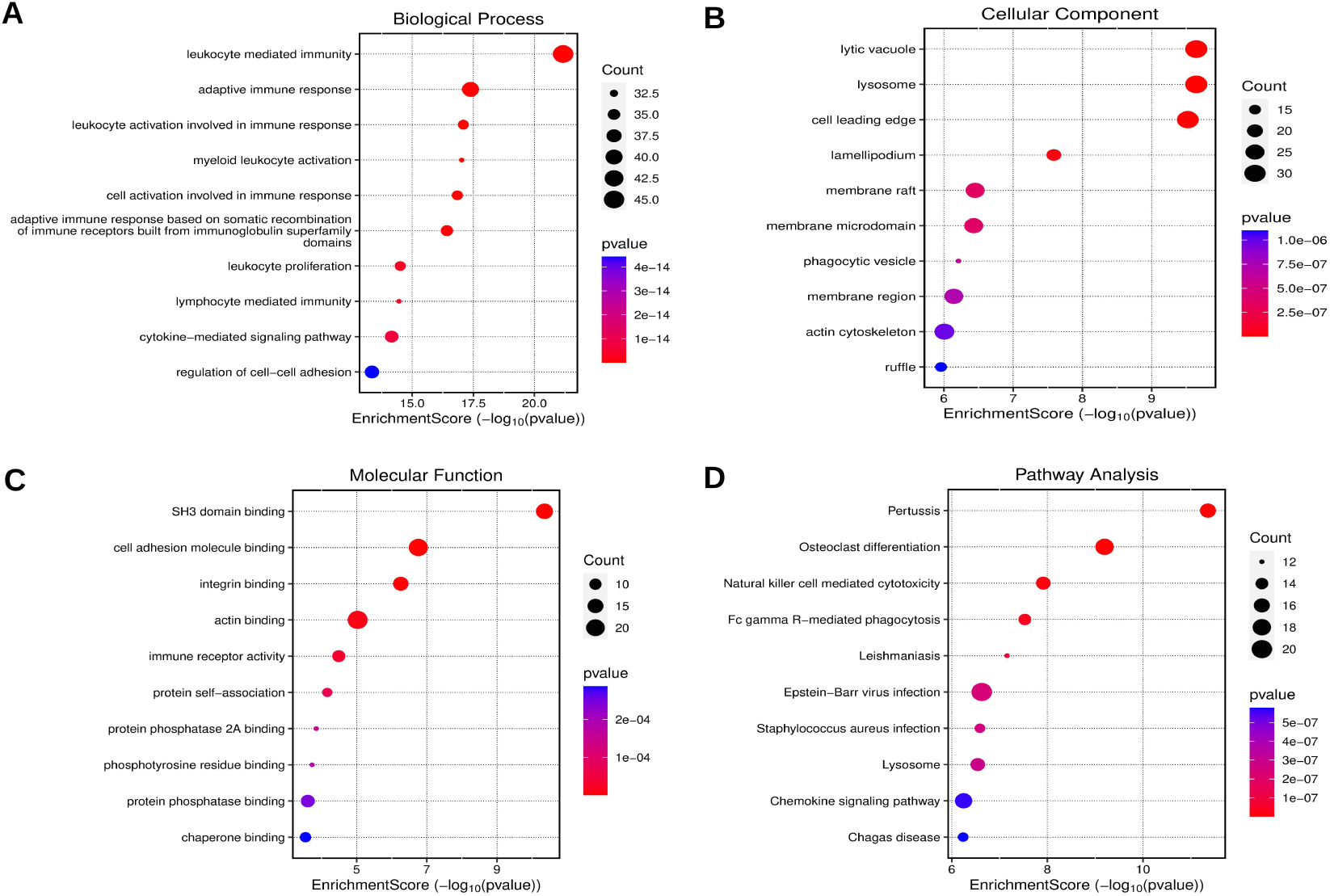
GO and KEGG enrichment analysis of TBI-related module genes using clusterProfiler. (A) The top enriched Biological process (B) The top enriched Cellular component (C) The top enriched Molecular function (D) The top enriched KEGG pathways.

### Protein Network and Hub Genes of T1DM and TBI

Using the STRING database, we built a PPI network of WGCNA T1DM-related module genes (**Fig. 10A**). Next, the cytoHubba plugin in Cytoscape was used to identify and visualize the hub genes within the network. In T1DM network, twenty hub genes: Rpl23, Rps3a, Hmgcs2, Rps6, Rpl5, Rpl17, Rps24, Rpl23a, Rps4x, Mtor, Pdk4, Rpl9, Rps15a14, Rpl30, Rpl31, Rps25, Rps27a-2, Kcna4, Slc2a4, and Cpt1a were selected based on the degree of their interaction in the network (**Fig. 10B**). Interestingly, the T1DM hub genes were divided into two clusters, ribosomal genes and mitochondrial genes (**Fig. 10B**). The ribosomal hub genes were primarily related to ribosome biogenesis and RNA post-transcriptional regulation, and function in rRNA binding. The mitochondrial genes were related to the lipid metabolic process and function in fatty-acyl-COA binding and insulin signaling. This highlights the key involvement of lipid metabolism and ribosome biogenesis in T1DM pathogenesis. Similarly using STRING, we constructed a PPI network of WGCNA TBI-related module genes (**Fig. 11A**). Next, using cytoHubba, twenty hub genes: Ptprc, Tp53, Stat1, Stat3, Tyrobp, Itgad, Csf1r, Itgb2, Rac2, Icam1, Myd88, Tgfb1, Vav1, Cd44, C1qa, Aif1, Lyn, B2m, Laptm5, and Fcer1g were selected based on the degree of their interaction in the network (**Fig. 11B**). The TBI hub genes were primarily related to inflammatory mediators and immune response. This points to the significant involvement of neuroinflammation in the pathophysiology of TBI and its complications.

**Fig. 10.**
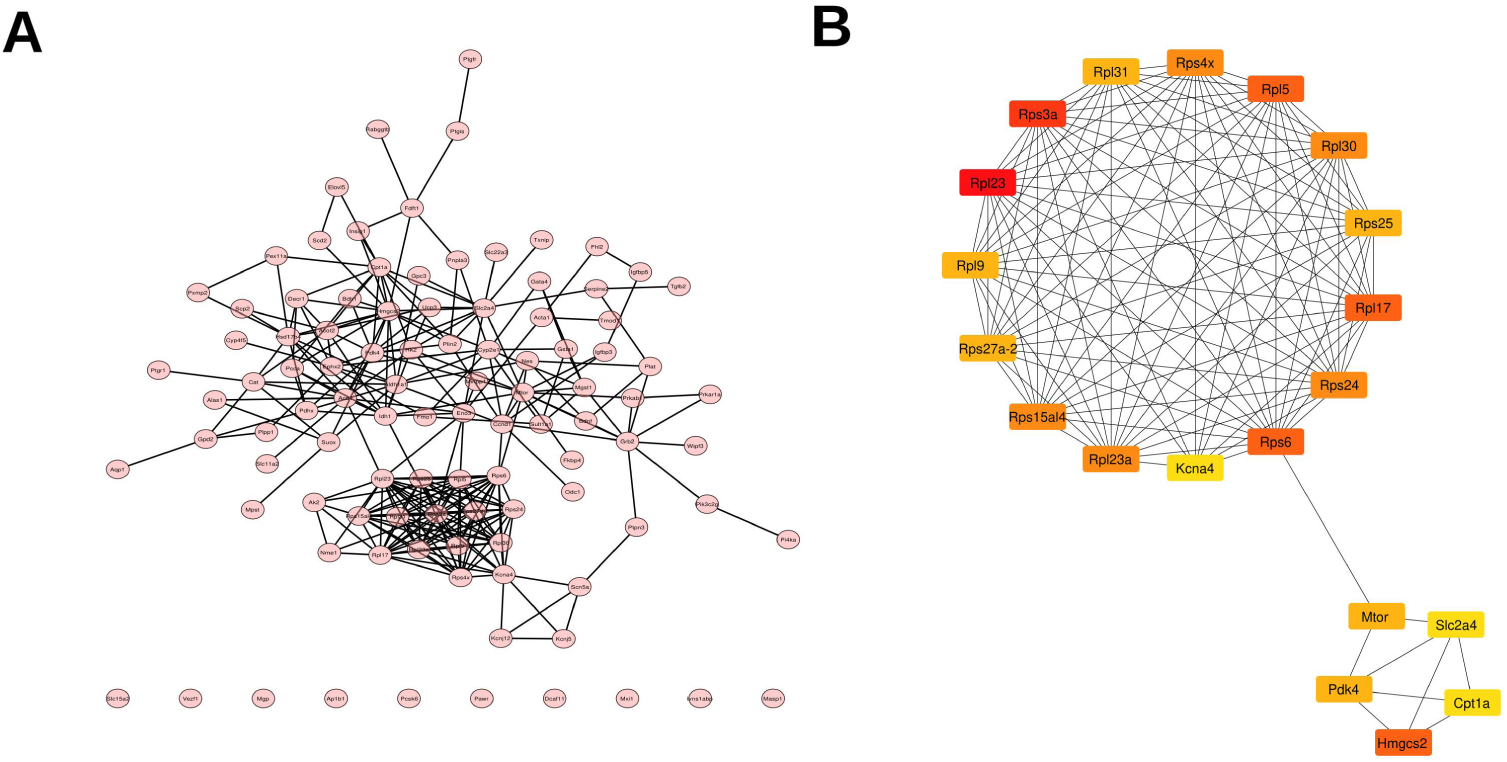
Protein Network analysis of T1DM-related module genes. (A) The PPI network diagram using confidence score of 0.4 constructed by STRING. (B) The top 20 hub genes selected based on the degree of interaction in the network. The red color means higher interaction and yellow color means lower interaction. STRING: Search Tool for the Retrieval of Interacting Genes/Proteins. PPI: Protein–protein interaction

**Fig. 11.**
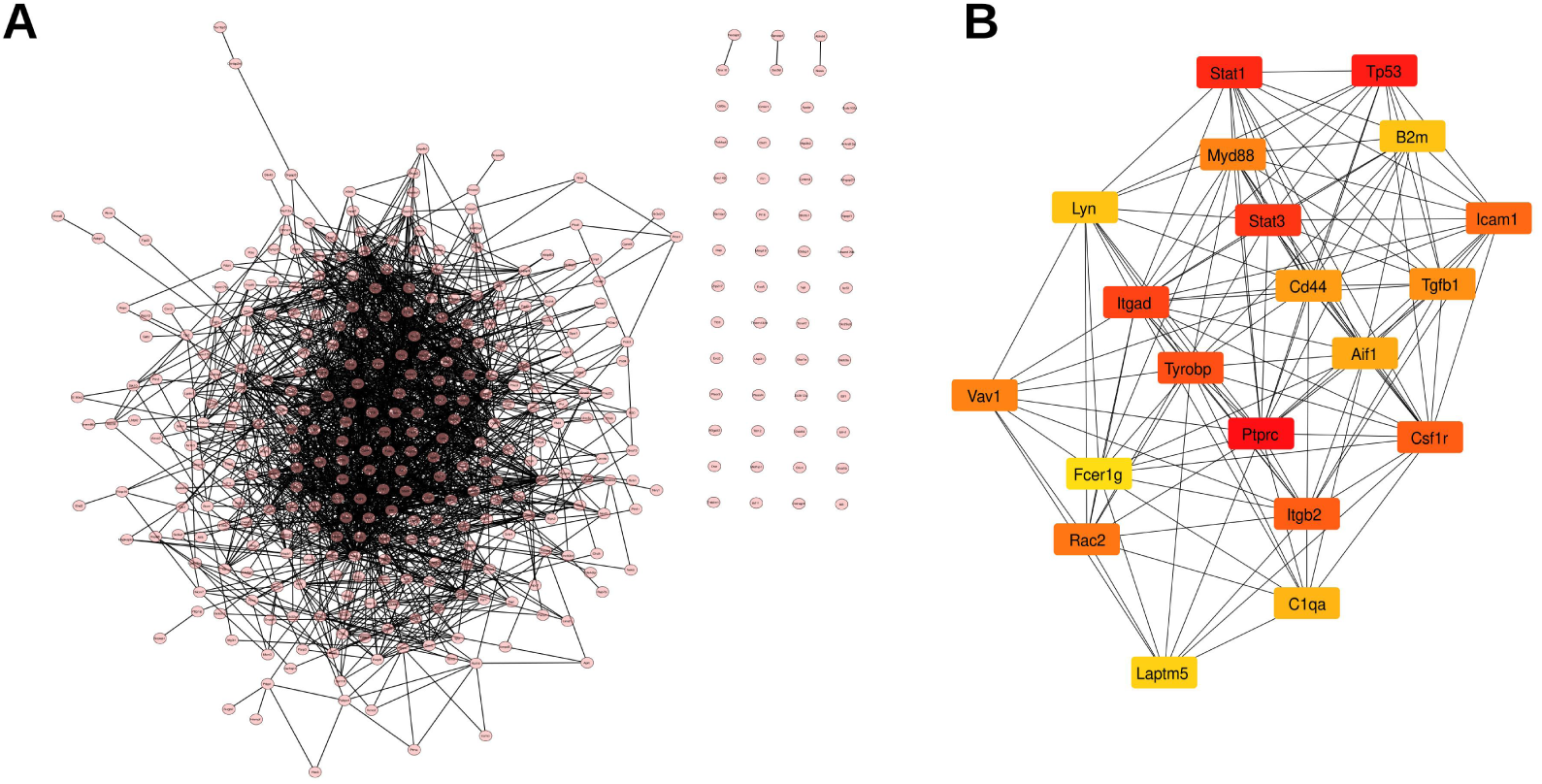
Protein Network analysis of TBI-related module genes. (A) The PPI network diagram using confidence score of 0.4 constructed by STRING. (B) The top 20 hub genes selected based on the degree of interaction in the network. The red color means higher interaction and yellow color means lower interaction. STRING: Search Tool for the Retrieval of Interacting Genes/Proteins. PPI: Protein–protein interaction

### Functional Enrichment Analysis of the Shared Genes of T1DM and TBI

The modules related to T1DM and TBI yielded a total of 122 and 368 genes respectively. A total of 3 shared genes were identified based on the overlap of genes related to T1DM and TBI (**Fig. 12A**), and these genes were considered to be highly correlated with the pathophysiology of T1DM-associated TBI. Next, we used “clusterProfiler” to examine the biological processes and enriched pathways to gain a better understanding of the functional roles of the shared genes in T1DM-associated TBI. Our findings indicated that the shared genes were enriched in biological processes such as cellular response to hydroperoxide, complement receptor-mediated signaling activity, negative regulation of IL-12 production, regulation of sequestering of triglyceride, positive regulation of lipid storage, and positive regulation of macrophage chemotaxis (**Fig. 12B**). The cine plot further shows the molecular functions specifically enriched by each of the shared genes (**Fig. 12C**).

**Fig. 12.**
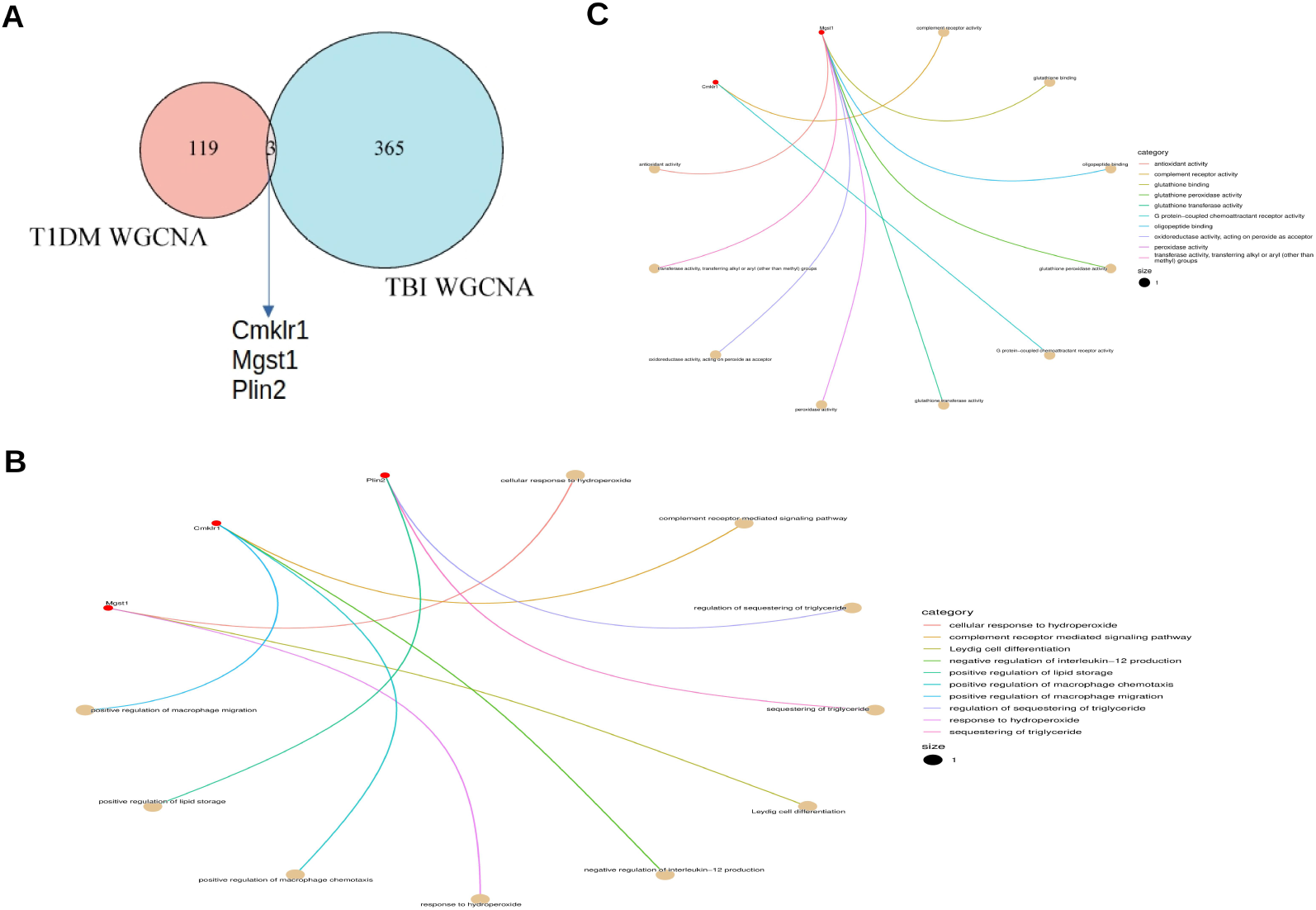
Functional enrichment and Identification of shared gene. (A) The Ven diagram of the shared genes. The genes were identified based on the overlap of TIDM-related module genes and TBI-related module genes. (B) The top enriched Biological process of the shared genes. (C) The top enriched Molecular function of the shared genes.

## DISCUSSION

Although there have been numerous studies on T1DM and TBI, the precise shared molecular mechanisms underlying these two conditions are still unknown. T1DM can coexist with TBI and recent studies have shown that diabetes mellitus exacerbates the outcomes of TBI, leading to more severe cognitive deficits and increased risk of complications [2]. In this study, we investigated the potential molecular mechanisms and identified key genes that are implicated in the pathogenesis of T1DM-associated TBI using WGCNA and bioinformatics analysis. Our findings indicate that neuroinflammation and the dysregulation of lipid metabolism represent two plausible molecular mechanisms underlying T1DM-associated TBI. We first identified key genes and significant pathways in each condition to get a better understanding of the biological processes that are involved in their pathogenesis.

### Dysregulation in Lipid metabolism and Oxidative stress implicated in Type 1 diabetes (T1MD)

We performed differential gene expression analysis to identify DEGs in T1DM dataset. The Top DEGs significantly upregulated are Acot2, Hmgcs2, Cpt1a, Txnip, Cbr1, and Pik3c2g. According to GSEA, these DEGs were significantly enriched in cell communication, lipid metabolic process, response to hypoxia, and PPAR signaling. The study of abnormalities in lipid metabolism has gained attention in an effort to better understand the pathogenesis of T1DM, even though altered glucose metabolism is still considered the primary driver of the disease. Our study revealed a significant upregulation in genes involved in lipid metabolic process. Acyl-CoA thioesterases (Acots) are hydrolases that regulate the balance between fatty acyl-CoAs and free fatty acids in cells [11]. Furthermore, Acots are regulated by PPAR agonists and nutritional factors, such as high fat feeding and fasting [11]. This highlights PPAR signaling enrichment in our analysis. Acot2 is a mitochondrial enzyme that hydrolyzes long-chain acyl-CoAs to produce free fatty acids and coenzyme A (CoASH), controlling their intracellular concentration. Acot2 has been linked to diabetes and is essential in controlling oxidative stress, inflammation, and hepatic fatty acid oxidation during fasting [12]. Hmgcs2 is a ketogenic enzyme that catalyzes the initial step in ketogenesis, that produces energy generated from lipid while fasting, and is implicated in T1DM [13]. Hmgcs2 was upregulated in T1DM and was implicated in T1DM-induced cardiac dysfunction [13]. Cpt1a is localized in the liver and is essential for fatty acid oxidation. Cpt1a help carnitine to bind long-chain fatty acids transporting them into the inner mitochondria membrane [14]. Long-chain fatty acids are a vital source of energy for the liver and other tissues during fasting time. Wei Sun et al. found that Cpt1a knockout mice showed a reduction in fasting insulin levels and a greater insulin glucose-lowering effect, indicating a greater disposal of glucose in the peripheral tissues [14]. The pathogenesis of T1DM hinges on the depletion of β-cells in the pancreas, which are accountable for the release of insulin. Txnip is one of the proteins that regulates the growth and degeneration of pancreatic β-cells [15]. Studies show that high Txnip levels cause β-cell death, while low Txnip prevents diabetes by increasing β-cell survival [15]. Thus, Txnip inhibitors could be promising therapeutic agents for managing diabetes related complications. Cbr1 is a key enzyme that detoxifies reactive aldehydes generated from lipid metabolism during oxidative stress [16]. One pathophysiology linked to pancreatic β-cell failure is oxidative stress [17]. Rashid et al. found that Cbr1 helps pancreatic β-cells survive glucotoxicity and glucolipotoxicity by shielding them from oxidative damage [17]. Pik3c2g is implicated in T2DM in humans [18]. Our analysis revealed an increase in Pik3c2g, but its precise role in T1DM remains unexplored. The PPI analysis of T1DM module genes identified 20 hub genes, including Hmgcs2, Cpt1a, Mtor, Pdk4, Kcna4, and Slc2a4, and 14 ribosomal genes, which will be discussed in the following section. The mTOR signaling pathway is implicated in metabolic syndrome, obesity, and diabetes [19], regulating glucose levels in the liver, adipose tissue, β-cells, and skeletal muscle through phosphorylation of IRS-1 [20]. It was found that rapamycin treatment of Mtor-activated mesangial cells reduced oxidative stress and apoptosis in cells exposed to high glucose, thereby improving diabetic nephropathy complications [21]. Pyruvate dehydrogenase complex (PDC) catalyzes the oxidative decarboxylation of pyruvate to acetyl-coenzyme A (CoA), which controls the entry of pyruvate into the TCA cycle [22]. This mechanism keeps the glucose-fatty acid cycle in check, maintaining glucose homeostasis [22]. However, Pdk4 inhibits PDC activity, and overexpression of Pdk4 is linked to metabolic disorders, such as diabetes and heart disease [23]. Thus, disrupting Pdk4 expression or using Pdk4 inhibitors may assist in treating these metabolic disorders. The Slc2a4 gene encodes the GLUT4 protein, which aids in glucose homeostasis in skeletal muscle and adipose tissue, suggesting increased expression of Slc2a4 could be a potential target for diabetes and insulin resistance [24]. Taken together, these studies suggest that disrupted lipid metabolism can cause oxidative stress and activate signaling pathways tied to T1DM complications. Hence, addressing aberrant changes in lipid metabolism could be an important strategy in mitigating diabetes complications.

### Ribosome biogenesis and RNA-binding proteins in the pathogenesis of Type 1 diabetes (T1DM)

We performed WGCNA on T1DM dataset and identified 122 significant genes that are positively correlated with T1DM. The PPI network analysis of these genes identified 20 hub genes that included 14 dominant ribosomal genes: Rpl23, Rps3a, Rps6, Rpl5, Rpl17, Rpl24, Rpl23a, Rps4x, Rpl9, Rps15a14, Rpl30, Rpl31, Rps25, and Rps27a-2. These genes were primarily related to ribosome biogenesis and RNA post-transcriptional regulation. These ribosomal genes were not detected by the limma analysis. One of the key benefits of WGCNA includes excellent sensitivity and system-level understanding of genes with low abundance or tiny fold changes [5]. Our analysis revealed that the ribosomal genes were localized in cytosolic ribosome and ribosomal subunit and function primarily in ribosome biogenesis and rRNA binding. Research on the specific function of ribosomal genes and their proteins in diabetes is limited, despite their extensive research in cancer [25]. This opens a new research avenue to delve deeper into the involvement of ribosomal genes in the pathogenesis of diabetes and its related complications. Interestingly, recent advancements in computational techniques and RNA-sequencing have uncovered irregular post-transcriptional gene regulation in diabetes involving ribosomal genes [26]. RNA-binding proteins (RBPs) play crucial role in controlling post-transcriptional RNA regulatory process, and have emerged as key players in diabetes and its related complications [26]. Dysregulation of RBPs is implicated in various diseases, including cancer, neurological, and cardiovascular disorders [26]. Recent studies have shown that RBPs contribute to the development of diabetes and diabetic complications through rRNA-binding mechanisms that include, alternative splicing (AS), RNA methylation, mRNA stability/degradation, and mRNA translation [26]. The topmost enriched RBP in our analysis is Rpl23. Rpl23 is a negative regulator of cellular apoptosis, and its overexpression is linked to aberrant apoptotic resistance in cancer [27]. Also, Rpl23 is implicated in diabetic retinopathy in T1DM C57BL/6 mice [28]. Similarly, Rps3a is elevated in T1DM ketoacidosis and positively correlates with activated CD56+CD16+ NK cells in this condition [29]. Additionally, Rps3a regulates mitochondrial function in human periaortic adipose tissue, causing atherosclerosis and vascular inflammation, which are diabetes-related complications [30]. Also, Rps6 is associated with prolonged hyperglycemia, reduced β-cell size, lowered insulin levels, and impaired glucose homeostasis [31]. Furthermore, Rpl24, Rpl31, and Rps3a were significantly upregulated in T1DM patients compared to healthy controls and increased the risk of developing T1DM in at-risk patients [32]. Similarly, elevated Rps4x in Turner syndrome (TS) significantly increases the risk of T1DM in women with X-chromosome monosomy [33]. A significant number of the identified ribosomal genes require further investigation in the context of T1DM to comprehend their specific roles. Taken together, ribosome biogenesis and RNA-binding proteins contribute to diabetes pathophysiology, but more research is needed to understand their specific roles in T1DM and target them for diabetes treatment.

### Microglia activation and Neuroinflammation implicated in Traumatic brain injury (TBI)

In the TBI dataset, we identified Vat1, Gfap, Tyrobp, Ctsz, Cd74, Ptpn6, Gpnmb, Itgad, and Itgb2 as the top significantly upregulated DEGs. In general, GSEA analysis showed that they were primarily enriched in the immune response-regulating signaling pathways. Specifically, metascape annotation showed enrichment of neutrophil activation and degranulation, positive regulation of tumor necrosis factor production, positive regulation of reactive oxygen species, Th1 and Th2 cell differentiation, Toll-like-receptor signaling pathway, and positive regulation of IL-6 production. The metascape annotation divided the DEGs into both negative and positive immune response mediators. Immune responses in TBI are now considered both damaging and beneficial, depending on the prevailing physiological condition [34]. Thus, if managed properly, the injured brain can also benefit from inflammation. The pathophysiology of TBI is divided into two separate phases: primary and secondary. Primary injury results from mechanical trauma to the head. During this acute phase of damage, neural tissue dies immediately and irreversibly [35]. In the initial acute phase of the injury, cytokines and chemokines activate microglia, triggering an immune response [36]. After TBI, inflammation is the primary driver of secondary brain damage, which can extend beyond the site of the lesion and continue over time, exacerbating the injury [36]. During initial injury, neuroinflammation is believed to be beneficial, but during secondary damage, persistent activation play a significant role in the progression of TBI [35]. The enrichment of DEGs in the signaling pathway for Toll-like receptors, Th1 and Th2 cell differentiation, and neurotrophil activation and degranulation suggests that these events contribute to neuroinflammation in TBI. In numerous experimental models of TBI, TLR-2 and TLR-4 have been demonstrated to play significant roles in inflammation and immune responses [37, 38]. Following TBI, mitochondrial release damage-associated molecular patterns (DAMPs), triggering production of cytokines, that activate toll-like receptors (TLRs) on cells and microglia [39]. DAMPs are also released by immune cells: monocytes, neutrophils and lymphocytes, at injury sites which exacerbate neuroinflammation [40]. Vat1 expression has been implicated in cancer, and Vat1-related genes were involved in immune and inflammatory responses [41]. The specific role of Vat1 in TBI is still unclear. Gfap is an intermediate filament protein produced by a variety of cell types, including astrocytes, and is used as a diagnostic biomarker for TBI. Elevated serum Gfap levels were seen within 1 hour after TBI, peaking 20 hours after the injury and then steadily declining 72 hours later [42]. Ctsz promotes chronic activation of the myeloid-derived microglia which exacerbates neuroinflammation in lipopolysaccharide (LPS) injected transgenic mice [43]. Cd74 has been shown to promote peripheral lymphocyte activation that exacerbates neurodegeneration in C57BL/6 TBI mice, and the inhibition of Cd74 decreased lymphocyte activation, lesion size, and neurodegeneration [44]. Gpnmb is highly expressed in macrophages and microglia, and has been associated with inflammation. However, the role of Gpnmb is still unclear as it is implicated in both proinflammatory and anti-inflammatory conditions [45]. Taken together, microglia activation and persistent neuroinflammation contribute to the secondary brain injury in TBI.

From the 368 TBI-related module genes, we identified 20 immune-related hub genes: Ptprc, Tp53, Stat1, Stat3, Tyrobp, Itgad, Csf1r, Itgb2, Rac2, Icam1, Myd88, Tgfb1, Vav1, Cd44, C1qa, Aif1, Lyn, B2m, Laptm5, and Fcer1g that are primarily associated with inflammation and immune responses. The topmost enriched genes were Ptprc, Tp53, Stat1, Stat3, Itgad, Itgb2 and Tyrobp. The topmost enriched biological processes associated with these genes were leukocyte-mediated immunity and adaptive immune response. Microglia activation, which generate inflammatory and cytotoxic factors, is the primary cause of persistent inflammation in most neurological disorders [46]. Activated microglia cause neuroinflammation by releasing cytotoxic and inflammatory molecules like IL-1β, TNF-α, and IL-6, exacerbating brain damage [47]. In a rat model of TBI, Ptprc was elevated at 24 hours but not at 4 hours after injury, promoting inflammation and enhancing NF-kappaB activity [48]. Plesnila et al. explored Tp53’s influence on neuron death, revealing it promotes neuronal apoptosis by hindering NF-kappaB. After TBI, p53 accumulates in injured brain cells, while NF-κB activity diminishes. Using pifithrin-α (PFT) to inhibit Tp53 dampens post-TBI neuron death, linked to elevated p53 levels [49]. Thus, Tp53 promotes neuronal apoptosis in damaged brains, making its inhibition a potential remedy for acute brain injuries. Stat1 promotes inflammation by activating proinflammatory gene expressions [50]. Suppressing IL-13 and the Jack1/Stat1 pathway inhibits pyroptosis, halting autophagy in moderate TBI [51]. On the other hand, Stat3 acts as an anti-inflammatory mediator by dampening cytokine signaling 3 and interleukins (IL-2 and IL-4) [52]. Itgad is mainly expressed on myeloid cells like macrophages, aiding leukocyte adhesion and activation. It was found that Itgad promotes brain inflammation and oxidative damage after TBI, and blocking it lessens brain inflammation and oxidative injury, improving neurological outcomes [53]. Also, elevated Tyrobp stimulates microglia to release cytokines, such as TNF-α, CXCL-8, and CCL2, which cause inflammation and neuronal apoptosis [54]. In contrast, Tyrobp knockout reversed activated microglia-induced inflammation and reduced neuronal cell apoptosis [54]. The study by Boone et al. indicates that the upregulation of the leukocyte-related gene Itgb2 persists even two weeks post-TBI injury, indicating its role in persistent inflammation activation [55]. Csf1r is a transmembrane tyrosine kinase receptor that is essential for microglia proliferation, differentiation, and survival [56]. Csf1r inhibition significantly reduced monocyte and neutrophil infiltration in the brain and blood at 1, 3, and 7 days post-TBI [57]. Rac2 is a member of the Rac family, which is expressed in neutrophils, macrophages, and adult T cells [58]. Rac2 facilitates microglial and astrocyte activation and promotes inflammation and apoptosis through the c-Jun N-terminal kinase (JNK) pathway in rats [58]. Rac2 deficiency reduces inflammation and oxidative stress and inhibits JNK to mitigate liver fibrosis [59]. However, the exact role of Rac2 in TBI remains unclear. Icam1 promotes adhesion and leukocyte infiltration of the brain [60]. Post-TBI, DAMPs stimulate microglia and endothelial cells to express Icam1 and release cytokines, promoting inflammation and neurodegeneration [60]. MyD88 signaling in hematopoietic cells and neutrophils contributes to inflammation after cortical brain damage, promoting the expression of proinflammatory cytokines IL-1β, IL-6, and CXCL16 [61]. Tgfb1 plays a dual role in inflammation, capable of both promoting and suppressing inflammatory responses, depending on the cellular environment and context in which it acts [62]. Vav1 is linked to microglial activation and inflammation in cerebral ischemia-reperfusion injury, and Vav1 knockout attenuated inflammation and apoptosis [63]. Furthermore, Vav1 is found to enhance NF-kB signaling, which intensifies inflammation in TBI [64]. Elevated Cd44 levels in TBI have been shown to exacerbate brain edema and neurological impairment [65]. C1qa is implicated in neuroinflammation and is associated with injury severity and poor prognosis following TBI [66]. Aif1 is expressed by activated microglia and infiltrating macrophages [67]. Aif1, Laptm5, Ptpn6, and Fcer1g were all upregulated in the hippocampus of TBI rats and were primarily associated with immune cell infiltration and inflammation [68]. Lyn is elevated in microglia and promotes Aβ plaque deposition in AD mice [69]. However, the role of Lyn in TBI is unclear. B2m is significantly elevated in patients with brain injury and is associated with impaired cognitive function due to its role in inflammation and immune response [70]. Taken together, these findings suggest that microglia activation and persistent inflammatory milieu are present within the injured brain following TBI. Therefore, the identified genes, along with the enriched biological processes, may elucidate the molecular mechanisms underlying TBI and offer potential therapeutic targets for TBI. Furthermore, our findings suggest that the modulation of inflammation-related factors and pathways constitutes a pivotal feature of TBI management.

### T1DM-associated TBI may involve Lipid droplet dysregulation and Neuroinflammation

In this study, we investigated the potential molecular mechanisms and identified key genes implicated in T1DM-associated TBI using WGCNA analysis. We found that Cmklr1, Mgst1, and Plin2 were concurrently present in the crucial modules linked to both T1DM and TBI, suggesting a high probability that they are involved in biological processes related to T1DM-associated TBI. Our analysis revealed that Cmklr1, Mgst1, and Plin2 were primarily enriched in biological processes related to lipid metabolism and immune response. The enriched biological processes were further corroborated by the GSEA conducted on T1DM and TBI datasets, indicating a strong link between the shared genes and the pathological mechanisms underlying T1DM-associated TBI. Furthermore, our analysis revealed that Cmklr1 regulates complement receptor signaling, IL-12 production, and macrophage chemotaxis and migration, while Plin2 regulates triglyceride sequestration and lipid storage, and Mgst1 is mainly involved in the cellular response to hydroperoxide. This suggests that T1DM-associated TBI may be linked to dysregulation of lipid metabolism and neuroinflammation.

Chemerin, an adipokine, regulates glycolipid metabolism and inflammation. It interacts with Cmklr1, a GPCR in innate immune cells. High levels of chemerin/Cmklr1 are linked to obesity and insulin resistance [71, 72]. Moreover, chemerin/Cmklr1 signaling axis is linked to diabetes-related complications like hypertension, myocardial infarction, heart failure, and cardiomyocyte apoptosis through inflammation [73, 74]. This suggests that chemerin/Cmklr1 plays a crucial role in diabetic cardiomyopathy by promoting cardiac inflammation. Also, chemerin/Cmklr1 is linked to diabetic nephropathy, but a study by Peng et al. found that α-NETA, a Cmklr1 antagonist, protects streptozotocin-induced mice from this damage [75]. Similarly, He and Mei’s clinical study on diabetic retinopathy (DR) revealed higher serum Cmklr1 levels in DR patients compared to healthy volunteers, and a positive correlation with diabetes duration, HbA1c, and LDL [76]. In a study by Tu et al. using C57BJ/6J mice fed with a high-fat diet, chemerin played a protective role in pancreatogenic diabetes mellitus (PDM). They found that chemerin levels decrease in patients with PDM, negatively affecting insulin resistance. Treatment with chemerin-9, a Cmklr1 agonist, increases chemerin levels, reducing glucose intolerance and insulin resistance in PDM model mice [77].

Taken together, chemerin/Cmklr1 signaling axis is implicated in the pathogenesis of diabetes and its related complications, including diabetic retinopathy, diabetic nephropathy, and diabetic cardiomyopathy. Also, chemerin/Cmklr1 signaling axis could have a protective role depending on the prevailing physiological condition, highlighting the dual function of Cmklr1. Our study revealed a positive correlation between Cmklr1 and T1DM, suggesting a potential role in T1DM pathogenesis, but further research is needed to understand its precise mechanism. It has also been found that Cmklr1 is crucial in chemerin-induced inflammation, acting as a strong chemoattractant to promote monocyte chemotaxis, thereby initiating inflammation [78]. However, depending on the ligands and physiological conditions, the Cmklr1 signaling pathway can have both pro- and anti-inflammatory effects [71]. This could explain the enrichment of complement receptor mediated signaling activity, negative regulation of IL-12 production and positive regulation of macrophage migration and chemotaxis by Cmklr1 in our analysis. In addition, our analysis revealed a positive correlation between Cmklr1 and TBI compared to healthy controls. Neuroinflammation is the predominant factor contributing to secondary brain injury in the aftermath of TBI [36]. The complement receptor signaling, negative regulation of IL-12 production and positive regulation of macrophage chemotaxis by Cmklr1 in our analysis could be attributed to its spatiotemporal role in modulating immune response [71]. It may lower IL-12 (anti-inflammatory) or promotes macrophage chemotaxis and migration (can be pro- or anti-inflammatory). Our explanation is that, Cmklr1 may initially contribute to inflammation by activating dendritic cells and macrophages, but later in inflammation, it may trigger pro-resolving pathways to mitigate it. Interestingly, Cmklr1 is implicated in neurological conditions like ischemic stroke (IS) and Alzheimer’s disease (AD) where it contributes to the polarization and migration of microglia [79, 80]. In AD mice, Chen et al. found that the chemerin/Cmklr1 axis is involved in the recruitment of microglia to Aβ deposition through the p38 MAPK pathway. In their work, Cmklr1 deficiency reduced the number of microglia around Aβ deposits in aged APP/PS1-CMKLR1−/− mice compared with APP/PS1 mice. Also, chemerin expression was significantly decreased in the hippocampus and cortex of aged APP/PS1 mice compared with WT mice, while the inhibition of p38 MAPK pathway attenuated microglial migration and polarization by chemerin/Cmklr1 signaling axis. They concluded that the chemerin/Cmklr1 axis is involved in the migration and recruitment of microglia to senile plaques via the p38 MAPK pathway, and modulation of the chemerin/Cmklr1 axis could be a potential new strategy for AD therapy [79]. In MCAO model of IS, Liu et al. found that Cmklr1 signaling ameliorates brain injury via inhibition of NLRP3 inflammasome-mediated neuronal pyroptosis [80]. They found that Cmklr1 expression was upregulated in MCAO, and Cmklr1 inhibition contributed to excessive NLRP3-mediated release of IL-1β and IL-18, as well as enhanced cleavage of GSDMD-N and neuronal pyroptosis, an indication of aggravated brain injury and neuronal damage. On the other hand, Cmklr1 activation by RvE1 or C-9 mitigated the neuronal damage, and overexpression of Cmklr1 in SH-SY5Y cells also rescued OGD-induced neuronal pyroptosis [80]. Previous studies have identified similarities in gene expression profiles and pathological mechanisms among AD, IS, and TBI [81, 82]. Thus, we believed that the Cmklr1 signaling axis might exert analogous effects in the context of TBI. However, whether these analogous effects are positive or negative in TBI remains unclear. From our analysis, Cmklr1 may contribute to the pathogenesis of T1DM and TBI by modulating inflammatory pathways crucial for both conditions. Additionally, Cmklr1’s role in diabetic complications like cardiomyopathy, retinopathy, and nephropathy may worsen TBI outcomes and increase the risk of complications. Therefore, involvement of Cmklr1 in inflammation could lead to antiinflammatory agents for T1DM-associated TBI, but its specific role in inflammation stages requires further investigation.

The second shared gene associated with T1DM and TBI we identified is Plin2 (perilipin 2). The perilipins (Plins) proteins are located on the periphery of lipid droplets (LDs) and play critical role in mediating intracellular lipid metabolism through the regulation of lipid droplet biogenesis and fatty acid sequestration [83]. Among Plins, Plin2 is the predominant isoform in nonadipocytes, particularly in β cells [84]. Also, Plin2 is implicated in disease progression of metabolic disorders and inflammation [83, 85]. Our analysis revealed that Plin2 is positively correlated with both T1DM and TBI, and is mainly enriched in the regulation of triglyceride sequestration, positive regulation of lipid storage, and positive regulation of macrophage chemotaxis. LDs are commonly used as location for sequestering and storing surplus lipid within cells [83]. Furthermore, LDs regulate β-cell function and β-cell death in diabetes [84]. Also, it was found that Plin2 is upregulated in pancreatic β-cells under nutritional stress [83]. Under physiological conditions, Plin2-coated LDs sequester fatty acids as triglycerides, averting lipid overload. This mechanism supports mitochondrial health and normal insulin secretion [83]. Under lipid overload, dietary stress leads to Plin2 downregulation in β-cells, impairing insulin production and damaging mitochondria [83]. Furthermore, when lipotoxicity triggers endoplasmic reticulum (ER) stress, β-cell failure occurs, leading to the shift from impaired glucose tolerance to diabetes [85]. Chen et al. discovered that diabetic Akita mice with a C96Y Ins2 mutation exhibit increased Plin2 expression and ER stress in their β-cells, but inhibiting Plin2 reduces stress, prevents β-cells death, and improves diabetes [85]. In summary, Plin2 expression increases in β-cells exposed to lipid loads or chemical ER stress, with Plin2 downregulation reducing ER stress effects, and overexpression exacerbates them [85]. Taken together, Plin2 deficiency impairs β-cell insulin production, leading to diabetes. From our analysis, Plin2 is positively correlated with T1DM, regulating triglyceride sequestration and lipid storage, suggesting its role in regulating abnormal lipid metabolism in T1DM. Therefore, targeting Plin2 to regulate lipotoxicity and ER stress may be a promising treatment approach for diabetes. There are limited studies on the expression of Plins in the human brain. In their study, Conte et al. found that Plins (Plin1 – Plin5) were expressed in various cerebral regions of the human brain [86]. They also conducted a correlation study with IL-6 expression to further investigate any potential associations between Plins and inflammation. They found that Plin2 was highly expressed in the gray matter of neurons, was elevated in aged individuals with AD pathology, and was positively correlated with IL-6 expression. They concluded that Plin2 is implicated in brain aging, and its accumulation could serve as a preliminary biomarker and a crucial precursor to inflammation and neurodegeneration [86]. Although brain aging and AD share many overlapping molecular mechanisms and disease pathology with TBI, the precise role of Plin2 in TBI remains unclear and warrants further investigation. Taken together, we infer that Plin2 may be involved in T1DM and TBI through the regulation of lipid sequestration and storage, and inflammation, and it could play a significant role in the molecular mechanisms underlying T1DM-associated TBI. It can also be a key therapeutic target for T1DM-associated TBI.

### Mgst1 (Microsomal Glutathione S-Transferase 1) may offer a protective mechanism for T1DM-associated TBI

The third shared gene linked to T1DM and TBI was identified as Mgst1, which is the most enriched of the three genes we identified. Mgst1 belongs to MAPEG (Membrane Associated Proteins in Eicosanoid and Glutathione metabolism), six family member proteins that are involved in eicosanoid and glutathione metabolism [87]. The eicosanoid family members are involved in the production of leukotrienes and prostaglandin E that mediate inflammation. The glutathione family members are involved in glutathione S-transferase and peroxidase activities and are collectively referred to as glutathione s-transferases (GSTs) [87, 88]. Mgst1 is one of the GSTs that are involved in cellular defense against toxic and carcinogenic substances, and represent major detoxification enzymes that protect cells from oxidative stress and inflammation [89, 90]. We found that Mgst1 is positively correlated with both T1DM and TBI and is primarily associated with the regulation of lipid hydroperoxides. When we looked at the molecular function of Mgst1, we found that Mgst1 mainly functions in pathways that respond to cellular stressors and protect cells from oxidative stress and inflammation. Thus, we postulate that Mgst1 may play a significant role in mitigating oxidative stress and inflammation, thereby enhancing the outcomes of T1DM-associated TBI.

Our study revealed that Mgst1 regulates cellular hydroperoxides and is primarily enriched in antioxidant activity, glutathione peroxidase activity, glutathione transferase activity, glutathione conjugation, and oxidoreductase activity (acting on peroxides as acceptor). Furthermore, our study revealed that Mgst1 is primarily localized in the peroxisome, microbody, and outer mitochondrial membranes, protecting them from oxidative stress and aiding in the detoxification of toxic and carcinogenic substances. Peroxisomes participate in oxidative processes, including lipid metabolism and catabolism of D-amino acids, polyamines, and bile acids, converting the reactive species and hydrogen peroxides produced in these processes into water using catalases and peroxidases, including Mgst1. Ferroptosis is a newly defined controlled cell death that is caused by iron-dependent lipid peroxidation. Recent studies have revealed that Mgst1 plays a crucial role in cellular processes and ferroptosis-related diseases, demonstrating its ability to mitigate their adverse effects. For example, Dai et al. found that the upregulation of Mgst1 can reduce the oxidative stress of trophoblast cells caused by hypoxia/reoxygenation [91]. Also, Mgst1 repressed ferroptosis-induced lipid peroxidation in pancreatic ductal adenocarcinoma (PDAC) by inactivating the lipid peroxide-generating enzyme arachidonate 5-lipoxygenase (ALOX5) [92]. Interestingly, ferroptosis is implicated in both diabetes and TBI [93, 94]. Recent studies indicate that ferroptosis significantly impacts the onset and progression of T1DM, T2DM, gestational diabetes, diabetic atherosclerosis, and various diabetes-related complications [93]. In TBI, ferroptosis induces secondary brain damage and promotes neuronal cell death and functional deficits, while the suppression of ferroptosis enhances recovery outcomes [94]. Thus, ferroptosis could be a novel therapeutic target for diabetes and TBI, and their related complications, and Mgst1 could be important in mitigating the detrimental effects of ferroptosis in T1DM-associated TBI.

Reactome was used to analyze the top enriched pathways of Mgst1, revealing its significance in modulating neutrophil degranulation and Phase II conjugation of electrophiles **(Fig. 13A and B**). Neutrophils play a crucial role in inflammation and are implicated in tissue injury across numerous pathological conditions, including brain injury [95]. Neutrophil degranulation refers to the secretion of pro-inflammatory factors by granulocytes within neutrophils [95] (**Fig. 13A**). In the innate immune response, neutrophils act as a double-edged sword, having both beneficial and destructive effects [95]. Following injury, neutrophils are rapidly recruited, infiltrating infarcts and interacting with necrosis and apoptosis cells to propagate inflammation, engulf dead cells, and facilitate reparative phase transformation [95]. However, excessive neutrophil degranulation drives inflammation by antigen presentation and secretion of cytokines, chemokines, leukotrienes, and prostaglandins, which cause tissue damage and worsen already-existing injury [96]. Furthermore, excessive neutrophil degranulation is a common feature of many inflammatory disorders, such as brain injury, ischemic stroke, acute lung injury, atherosclerosis, and rheumatoid arthritis [96, 97]. Thus, we believe Mgst1 may play a role in regulating the inflammatory response and neutrophil extracellular trap formation in T1DM-associated TBI by regulating the exocytosis of azurophil granule lumen proteins. Therefore, the modulation of neutrophil degranulation by Mgst1 could be an important therapeutic strategy for T1DM-associated TBI. Furthermore, Mgst1 is involved in the biotransformation reaction through phase II conjugation of electrophiles using glutathione [98] Metabolism of xenobiotics produce electrophilic or nucleophilic compounds with reactive groups such as epoxides, carbonyl, hydroxyls, amino and sulfhydryl groups capable of attacking electron-rich DNA and proteins [99]. In addition, these reactive oxidants are generated endogenously (e.g. from oxidative phosphorylation or phagocytosis), and if accumulate results in oxidative stress and inflammation [99]. These reactive groups are conjugated with glucuronide, sulfate, acetate or amino acids and transformed into hydrophilic molecules, which are easily eliminated by bile and urine [99] (**Fig. 12B**). The conjugation and biotransformation reactions are catalyze by glutathione S-transferases (Mgst1 and other GSTs) and other phase II conjugation enzymes (**Fig. 13B**). Taken together, oxidative stress and inflammation are key pathological features of T1DM and TBI, and Mgst1 can alleviate these detrimental effects by modulating the secretion of inflammatory factors from neutrophils and facilitate phase II conjugation of electrophiles to promote their elimination, thereby providing a protective mechanism against T1DM-associated TBI.

**Fig. 13.**
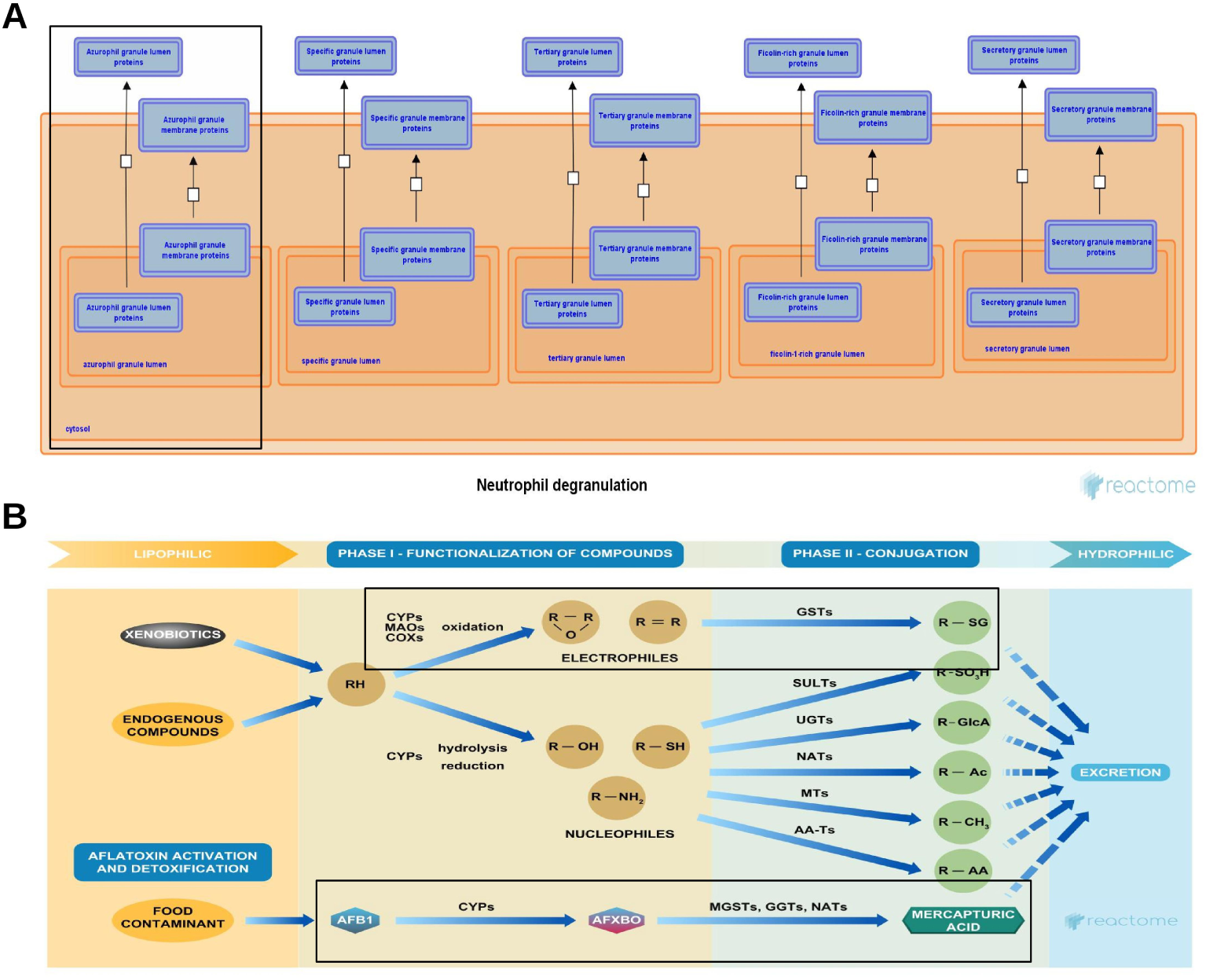
Reactome pathway diagram. (A) The role of Mgst1 in the modulation of Neutrophil degranulation. (B) The role of Mgst1 in Phase II conjugation of electrophiles. Mgst1 – Microsomal Glutathione S-Transferase 1. Pathway diagram provided by Reactome Database (www.reactome.org) under Creative Commons Attribution 4.0 License.

## CONCLUSION

Overall, our findings identified key genes and enriched biological processes in T1DM-associated TBI. We revealed that genes related to the lipid metabolic pathway and ribosome biogenesis are involved in T1DM pathogenesis. Ribosome biogenesis and RNA-binding proteins are emerging as key players in diabetes and its related complications. Furthermore, genes identified in TBI were primarily related to inflammation and immune response, highlighting the significant involvement of neuroinflammation in TBI pathogenesis. Our study, for the first time, identified Cmklr1, Mgst1, and Plin2 as key genes of T1DM-associated TBI, and they may play important roles in the molecular mechanisms of T1DM-associated TBI via regulating lipid droplet sequestration and inflammation. However, the current study has some limitations. First, the sample size of the GEO datasets is relatively small, and future studies might require larger sample sizes for experiment and validations. Second, we investigated primarily the molecular mechanism between Type 1 diabetes and TBI, and the findings might not provide sufficient evidence for Type II diabetes. Third, the findings in the current study are based on bioinformatics analysis, and the results lacked the validation of in-vivo experiments. Finally, more research and validations are needed to determine whether Cmklr1, Mgst1, and Plin2 have clinical diagnostic and therapeutic relevance for T1DM-associated TBI.

## Author Contributions

Conceptualization and study design: UFS; Acquisition, assembly, and preprocessing of data; UFS, IB; Performed the computational analysis: UFS; Result interpretation and visualization: UFS; Writing of the manuscript: UFS. All authors read and approved the final manuscript.

## Data availability statement

The datasets used in this study are freely accessible and can be downloaded from the Gene Expression Omnibus (GEO) database at https://www.ncbi.nim.nih.gov/gds/ using their accession number GSE4745, GSE125451, GSE173975, and GSE80174.

## Supplementary material

For reproducibility, the R-markdown scripts for the study can be accessed on the GitHub repository https://github.com/UFSaidu/WGCNA_of_T1DM_associated_TBI.

## Acknowledgements

We acknowledge the Gene Expression Omnibus (GEO) database for providing free access to their platforms and the data contributors for uploading their transcriptome datasets.

---

**Table S1:** A Summary of GEO datasets used in the current study.

| **Dataset** | **Model** | **Condition** | **Tissue** | **Organism** | **Platform** | **Total samples**<br>**(disease/control)** |
| --- | --- | --- | --- | --- | --- | --- |
| GSE4745 | Streptozotocin | Diabetes | Heart | Rattus<br>norvegicus | GPL85 | 24 (12/12) |
| GSE125451 | Streptozotocin | Diabetes | Heart | Rattus<br>norvegicus | GPL20084 | 16 (12/4) |
| GSE173975 | Lateral fluid<br>percussion | Traumatic<br>brain injury | Hippocampus | Rattus<br>norvegicus | GPL22396 | 22 (11/11) |
| GSE80174 | Lateral fluid<br>percussion | Traumatic<br>brain injury | Hippocampus | Rattus<br>norvegicus | GPL14844 | 10 (5/5) |

